# An Indexing Theory for Working Memory based on Fast Hebbian Plasticity

**DOI:** 10.1101/334821

**Authors:** Florian Fiebig, Pawel Herman, Anders Lansner

**Affiliations:** Lansner Laboratory, Department of Computational Science and Technology, Royal Institute of Technology, 10044 Stockholm, Sweden; Department of Mathematics, Stockholm University, 10691 Stockholm, Sweden

## Abstract

Working memory (WM) is a key component of human memory and cognition. Computational models have been used to study the underlying neural mechanisms, but neglected the important role of short- and long-term memory interactions (STM, LTM) for WM. Here, we investigate these using a novel multi-area spiking neural network model of prefrontal cortex (PFC) and two parieto-temporal cortical areas based on macaque data. We propose a WM indexing theory that explains how PFC could associate, maintain and update multi-modal LTM representations. Our simulations demonstrate how simultaneous, brief multi-modal memory cues could build a temporary joint memory representation as an “index” in PFC by means of fast Hebbian synaptic plasticity. This index can then reactivate spontaneously and thereby reactivate the associated LTM representations. Cueing one LTM item rapidly pattern-completes the associated un-cued item via PFC. The PFC-STM network updates flexibly as new stimuli arrive thereby gradually over-writing older representations.

## Introduction

By working memory (WM), we typically understand a flexible but volatile kind of memory capable of holding a small number of items over short time spans, allowing us to act beyond the immediate here and now. WM is thus a key component in cognition and is often affected early on in neurological and psychiatric conditions, e.g. Alzheimer’s disease and schizophrenia (Slifstein et al. 2015). Prefrontal cortex (PFC) has consistently been implicated as a key neural substrate for WM in humans and non-human primates (Fuster 2009; D’Esposito & Postle 2015).

Computational models of WM have so far focused mainly on its short-term memory aspects, explained either by means of persistent activity (Funahashi et al. 1989; Goldman-Rakic 1995; Camperi & Wang 1998; Compte et al. 2000) or more recently fast synaptic plasticity (Mongillo et al. 2008; Fiebig & Lansner 2017; Lundqvist et al. 2011) as the underlying neural mechanism. The nature of neural mechanisms involved in WM processes in PFC and, consequently, their neural manifestations have strong implications for the dynamic interaction between short- and long-term memory (STM, LTM). Although this operational STM-LTM coupling has been often missing in computational and empirical studies, it is critical to WM function as it enables to activate or “bring online” a small set of WM task relevant LTM representations (Eriksson et al. 2015). This prominent effect is envisaged to underlie complex cognitive phenomena, which have been characterized extensively in experiments on humans as well as animals. Nevertheless, the neural mechanisms involved remain elusive.

Here we present a large-scale spiking neural network model of WM and focus on investigating the neural mechanisms behind the fundamental STM-LTM interactions critical to WM function. In this context, we introduce a WM indexing theory, inspired by the predecessor hippocampal memory indexing theory (Teyler & DiScenna 1986) originally proposed to account for the role of hippocampus in storing episodic memories (Teyler & Rudy 2007). Notably, Teyler and Rudy (2007) emphasized the role of rapid hippocampal synaptic plasticity for indexing to work. We propose that Hebbian plasticity in PFC could be even faster and serve as a key mechanism in synaptic WM. The phenomena of binding and indexing of neural representations have been a common recurring theme in memory research, in particular in relation to the role of hippocampus and surrounding structures (Teyler & Rudy 2007; Squire 1992; O’Reilly & Frank 2006). Our main novel contribution here is to show that a neurobiologically constrained large-scale spiking neural network model of interacting cortical areas can function as a robust WM, including its important role of bringing relevant LTM representations temporarily on-line by means of “indexing”. In addition, the model replicates many experimentally observed phenomena in terms of oscillations, coherence and latency within and between cortical regions.

The core idea of our theory rests on the concept of cell assemblies formed in the PFC by means of fast Hebbian plasticity that serve as “indices” linking LTM representations. Our model comprises a subsampled PFC network model of STM that is reciprocally connected with two LTM component networks representing different sensory modalities (e.g. visual and auditory) in temporal cortical areas. This new model builds on and extends our recent PFC-dependent STM model of human word-list learning (Fiebig & Lansner 2017), shown to reproduce a range of patterns of mesoscopic neural activity observed in WM experiments, and it employs the same fast Hebbian plasticity as a key neural mechanism, intrinsically within PFC but also in PFC backprojections that target parieto-temporal LTM stores. This novel concept, at the heart of our WM indexing theory, has strong implications for WM function and results in large-scale inter-network dynamics as a neural correlate of WM phenomena, which offers macroscopic predictions for experimental validation. Plasticity in this functional context needs to be Hebbian, i.e. associative, and has to be induced and expressed on a time-scale of a few hundred milliseconds. Recent experiments have demonstrated the existence of fast forms of Hebbian synaptic plasticity, e.g. short-term potentiation (STP, or Labile LTP) (Erickson et al. 2010; Park et al. 2014; Kauer et al. 2018), which lends credibility to this type of WM mechanism.

We hypothesize that activity in parieto-temporal LTM stores targeting PFC via fixed patchy synaptic connections triggers an activity pattern in PFC, which is rapidly connected by means of fast Hebbian plasticity to form a cell assembly displaying attractor dynamics. The connections in backprojections from PFC to the same LTM stores are also enhanced and provide a functional link specifically with the triggering cell assemblies there. Our simulations demonstrate that such a composite WM model can function as a robust and flexible multi-item and cross-modal WM that maintains a small set of activated task relevant LTM representations and associations. Transiently formed cell assemblies in PFC serve the role of indexing and temporary binding of these LTM representations, hence giving rise to the name of the proposed theory. The PFC cell assemblies can activate spontaneously and thereby reactivate the associated long-term representations. Cueing one LTM item rapidly activates the associated un-cued item via PFC by means of pattern completion. The STM network flexibly updates WM content as new stimuli arrive whereby older representations gradually fade away. Interestingly, this model implementing the WM indexing theory can also explain the so far poorly understood cognitive phenomenon of variable binding or object – name association, which is one key ingredient in human reasoning and planning (Cer & O’Reily 2012; van der Velde & de Kamps 2015; Pinkas et al. 2013).

## Materials & Methods

### Neuron Model

We use an integrate-and-fire point neuron model with spike-frequency adaptation (Brette & Gerstner 2005) which was modified (Tully et al. 2014) for compatibility with a custom-made BCPNN synapse model in NEST (see *Simulation Environment*) through the addition of the intrinsic excitability current 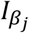. The model was simplified by excluding the subthreshold adaptation dynamics. Membrane potential *V*_*m*_ and adaptation current are described by the following equations:

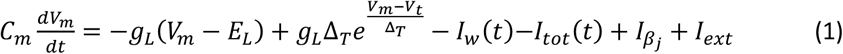

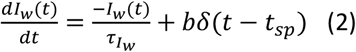

The membrane voltage changes through incoming currents over the membrane capacitance *C*_*m*_. A leak reversal potential *E*_*L*_ drives a leak current through the conductance *g*_*L*_, and an upstroke slope factor Δ_*T*_ determines the sharpness of the spike threshold *V*_*t*_. Spikes are followed by a reset of membrane potential to *V*_*r*_. Each spike increments the adaptation current by *b*, which decays with time constant 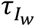. Simulated basket cells feature neither the intrinsic excitability current 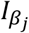 nor this spike-triggered adaptation.

Besides external input *I*_*ext*_ (*Stimulation Protocol*) neurons receive a number of different synaptic currents from its presynaptic neurons in the network (AMPA, NMDA and GABA), which are summed at the membrane accordingly:

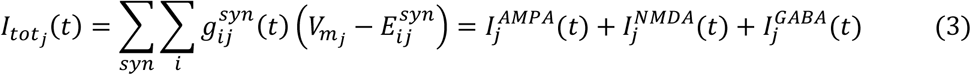

### Synapse Model

Excitatory AMPA and NMDA synapses have a reversal potential *E*^*AMPA*^ = *E*^*NMDA*^, while inhibitory synapses drive the membrane potential toward *E*^*GABA*^. Every presynaptic input spike (at 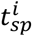 with transmission delay *t*_*ij*_) evokes a transient synaptic current through a change in synaptic conductance that follows an exponential decay with time constants *τ*^*syn*^ depending on the synapse type (*τ*^*AMPA*^ ≪ *τ*^*NMDA*^).

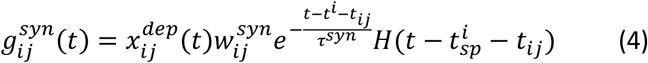

*H*(·) is the Heaviside step function. 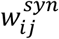 is the peak amplitude of the conductance transient, learned by the *Spike-based BCPNN Learning Rule* (next Section). Plastic synapses are also subject to synaptic depression (vesicle depletion) according to the Tsodyks-Markram formalism (Tsodyks & Markram 1997), modeling the transmission-dependent depletion of available synaptic resources 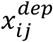 by a utilization factor U, and a depression/reuptake time constant *τ*_*rec*_:

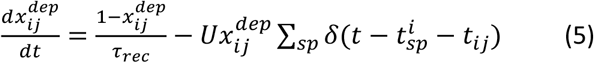

### Spike-based BCPNN Learning Rule

Plastic AMPA and NMDA synapses are modeled to mimic short-term potentiation (STP) (Erickson et al. 2010) with a spike-based version of the Bayesian Confidence Propagation Neural Network (BCPNN) learning rule (Wahlgren & Lansner 2001; Tully et al. 2014). For a full derivation from Bayes rule, deeper biological motivation, and proof of concept, see Tully et al. (2014) and the earlier STM model implementation (Fiebig & Lansner 2017).

Briefly, the BCPNN learning rule makes use of biophysically plausible local traces to estimate normalized pre- and post-synaptic firing rates, as well as co-activation, which can be combined to implement Bayesian inference because connection strengths and neural unit activations have a statistical interpretation (Sandberg et al. 2002; Fiebig & Lansner 2014; Tully et al. 2014). Crucial parameters include the synaptic activation trace Z, which is computed from spike trains via pre- and post-synaptic time constants 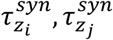, which are the same here but differ between AMPA and NMDA synapses:

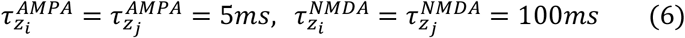

The larger NMDA time constant reflects the slower closing dynamics of NMDA-receptor gated channels. All excitatory connections are drawn as AMPA and NMDA pairs, such that they feature both components. Further filtering of the Z traces leads to rapidly expressing memory traces (referred to as P-traces) that estimate activation and coactivation:

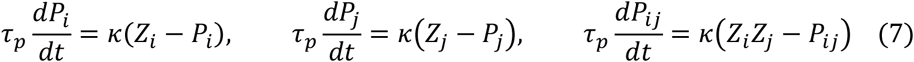

These traces constitute memory itself and decay in a palimpsest fashion. STP decay is known to take place on timescales that are highly variable and activity dependent (Volianskis et al. 2015); see Discussion – The case for Hebbian plasticity.

We make use of the learning rule parameter *κ* (Equation 7), which may reflect the action of endogenous neuromodulators, e.g. dopamine acting on D1 receptors that signal relevance and thus modulate learning efficacy. It can be dynamically modulated to switch off learning to fixate the network, or temporarily increase plasticity (*κ*_*p*_, *κ*_*normal*_, Table 1). In particular, we trigger a transient increase of plasticity concurrent with external stimulation.

Tully et al. (2014) showed that Bayesian inference can be recast and implemented in a network using the spike-based BCPNN learning rule. Prior activation levels are realized as an intrinsic excitability of each postsynaptic neuron, which is derived from the post-synaptic firing rate estimate p_j_ and implemented in the NEST neural simulator (Gewaltig & Diesmann 2007) as an individual neural current 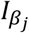 with scaling constant β_gain_

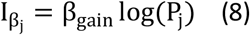

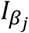 is thus an activity-dependent intrinsic membrane current to the neurons, similar to the A-type potassium channel (Hoffman et al. 1997) or TRP channel (Petersson et al. 2011). Synaptic weights are modeled as peak amplitudes of the conductance transient (Equation 4) and determined from the logarithmic BCPNN weight, as derived from the P-traces with a synaptic scaling constant 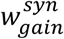.

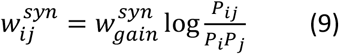

In our model, AMPA and NMDA synapses make use of 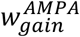 and 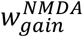 respectively. The logarithm in Equations 8, 9 is motivated by the Bayesian underpinnings of the learning rule, and means that synaptic weights 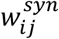 multiplex both the learning of excitatory and di-synaptic inhibitory interaction. The positive weight component is here interpreted as the conductance of a monosynaptic excitatory pyramidal to pyramidal synapse (Figure 1, plastic connection to the co-activated MC), while the negative component (Figure 1, plastic connection to the competing MC) is interpreted as di-synaptic via a dendritic targeting and vertically projecting inhibitory interneuron like a double bouquet and/or bipolar cell (Tucker 2002; Ren et al. 2007; Silberberg & Markram 2007; Kapfer et al. 2007). Accordingly, BCPNN connections with a negative weight use a GABAergic reversal potential instead, as in previously published models (Tully et al. 2016; Tully et al. 2014; Fiebig & Lansner 2017). Model networks with negative synaptic weights have been shown to be functionally equivalent to ones with both excitatory and inhibitory neurons with only positive weights (Parisien et al. 2008).

Code for the NEST implementation of the BCPNN synapse is openly available (see *Simulation Environment*).

**Figure 1.**
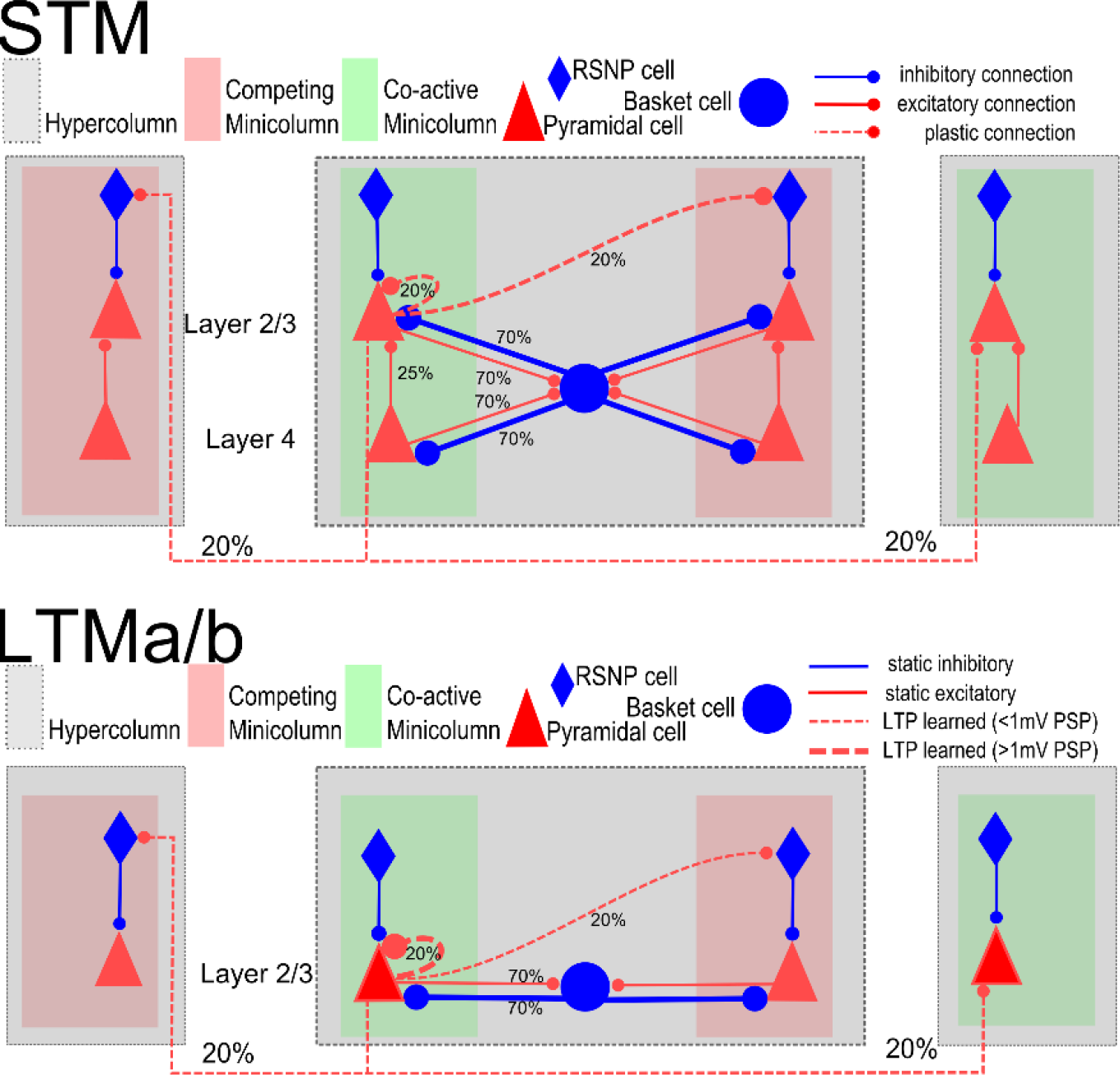
Local columnar connectivity within STM and LTM. Connection probabilities are given by the percentages, further details in Tables 1–3. The strength of plastic connections develops according to the synaptic learning rule described in *Spike-based BCPNN Learning Rule*. Initial weights are low and distributed by a noise-based initialization procedure (*Stimulation protocol*). LTM however, dashed connections are not plastic in LTM (besides the STD of Equation 4), but already encode memory patterns previously learned through an LTP protocol, and loaded before the simulation using receptor-specific weights found in Table 2.

### Axonal Conduction Delays

We compute axonal delays *t*_*ij*_ between presynaptic neuron i and postsynaptic neuron j, based on a constant conduction velocity *V* and the Euclidean distance between respective columns. Conduction delays were randomly drawn from a normal distribution with mean according to the connection distance divided by conduction speed and with a relative standard deviation of 15% of the mean in order to account for individual arborization differences. Further, we add a minimal conduction delay 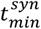 of 1.5 ms to reflect not directly modeled delays, such as diffusion of transmitter over the synaptic cleft, dendritic branching, thickness of the cortical sheet, and the spatial extent of columns:

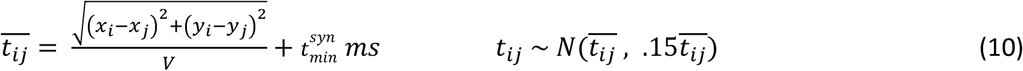

### STM Network Architecture

The model organizes cells in the three simulated cortical areas into grids of nested hypercolumns (HCs) and minicolumns (MCs), sometimes referred to as macro columns, and “functional columns” respectively. The STM network is simulated with 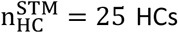 spread out on a grid with spatial extent of 17×17 mm. This spatially distributed network of columns has sizable conduction delays due to the distance between columns and can be interpreted as a spatially distributed subsampling of columns from the extent of dorsolateral PFC (such as BA 46 and 9/46, which also have a combined spatial extent of about 289 mm^2^ in macaque).

Each of the non-overlapping HCs has a diameter of about 640 μm, comparable to estimates of cortical column size (Mountcastle 1997), contains 48 basket cells, and its pyramidal cell population has been divided into twelve MC’s. This constitutes another sub-sampling from the roughly 100 MC per HC when mapping the model to biological cortex. We simulate 20 pyramidal neurons per MC to represent roughly the layer 2 population of an MC, 5 cells for layer 3A, 5 cells for layer 3B, and another 30 pyramidal cells for layer 4, as macaque BA 46 and 9/46 have a well-developed granular layer (Petrides & Pandya 1999). The STM model thus contains about 18,000 simulated pyramidal cells in four layers (although layers 2, 3A, and 3B are often treated as one layer 2/3).

**Figure 2.**
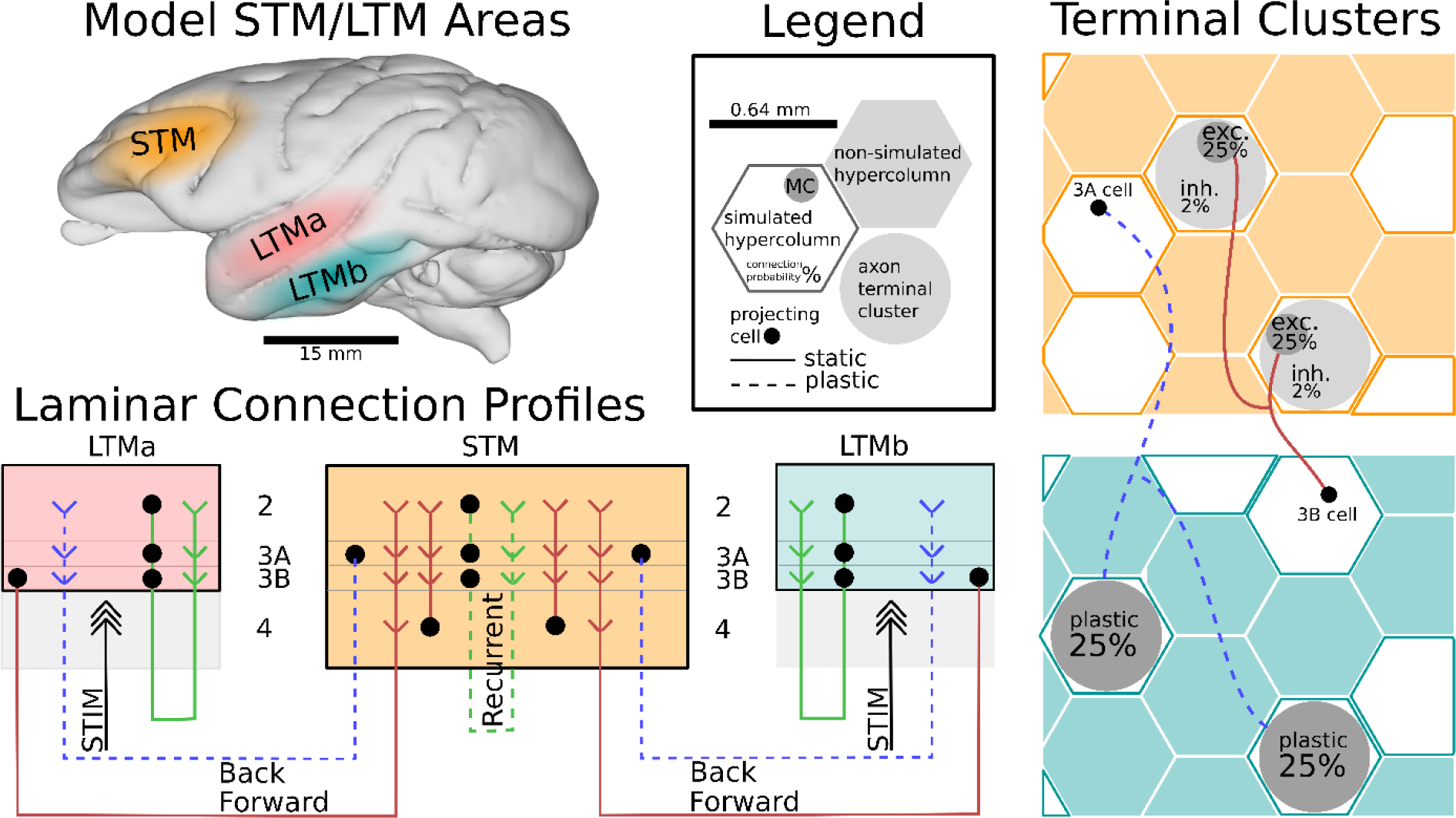
Schematic of modeled connectivity within and across representative STM and LTM areas in macaque. STM features 25 hypercolumns (HC), whereas LTMa and LTMb both contain 16 simulated HCs. Each network spans several hundred mm^2^ and the simulated columns constitute a spatially distributed subsample of biological cortex, defined by conduction delays. Pyramidal cells in the simulated supragranular layers form connections both within and across columns. STM features an input layer 4 that shapes the input response of cortical columns, whereas LTM is instead stimulated directly to cue the activation of previously learned long-term memories. Additional corticocortical connections (feedforward in brown, feedback in dashed blue) are sparse (<1% connection probability) and implemented with terminal clusters (rightmost panels) and specific laminar connection profiles (bottom left). The connection schematic illustrates laminar connections realizing a direct supragranular forward-projection, as well as a common supragranular backprojection. Layer 2/3 recurrent connections in STM (dashed green) and corticocortical backprojections (dashed blue) feature fast Hebbian plasticity. For an in-depth model description, including the columnar microcircuits, please refer to *Methods* and Figure 1.

### STM Network Connectivity

The most relevant connectivity parameters are found in Tables 1–3. Pyramidal cells project laterally to basket cells within their own HC via AMPA-mediated excitatory projections with a connection probability of *p*_*P–B*_, i.e. connections are randomly drawn without duplicates until the target fraction of all possible pre-post connections exist. In turn, they receive GABAergic feedback inhibition from basket cells (*p*_*B–P*_) that connect via static inhibitory synapses rather than plastic BCPNN synapses. This strong loop implements a competitive soft-WTA subnetwork within each HC (Douglas & Martin 2004). Local basket cells fire in rapid bursts, and induce alpha/beta oscillations in the absence of attractor activity and gamma, when attractors are present and active.

Pyramidal cells in layer 2/3 form connections both within and across HCs at connection probability *p*_*L23e–L23e*_. These projections are implemented with plastic synapses and contain both AMPA and NMDA components, as explained in subsection *Spike-based BCPNN Learning Rule*. Connections across columns and areas may feature sizable conduction delays due to the implied spatial distance between them (Table 1)

Pyramidal cells in layer 4 project to pyramidal cells of layer 2/3, targeting 25% of cells within their respective MC only. Experimental characterization of excitatory connections from layer 4 to layer 2/3 pyramidal cells have confirmed similarly high fine-scale specificity in rodent cortex (Yoshimura & Callaway 2005) and in-turn, full-scale cortical simulation models without functional columns have found it necessary to specifically strengthen these connections to achieve defensible firing rates (Potjans & Diesmann 2014).

In summary, the STM model thus features a total of 16.2 million plastic AMPA- and NMDA-mediated connections between its 18,000 simulated pyramidal cells, as well as 67,500 static connections from 9,000 layer 4 pyramidals to layer 2/3 targets within their respective MC, and 1.2 million static connections to and from 1,200 simulated basket cells.

### LTM network

We simulate two structurally identical LTM networks, referred to as LTMa, and LTMb. LTM networks may be interpreted as a spatially distributed subsampling of columns from areas of the parieto-temporal cortex commonly associated with modal LTM stores. For example Inferior Temporal Cortex (ITC) is often referred to as the storehouse of visual LTM (Miyashita 1993). Two such LTM areas are indicated in Figure 2.

We simulate 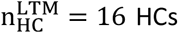 in each area and nine MC per HC (further details in Tables 1–3). Both LTM networks are structurally very similar to the previously described STM, yet they do not feature plasticity among their own cells, beyond short-term dynamics in the form of synaptic depression. Unlike STM, LTM areas also do not feature an input layer 4, but are instead stimulated directly to cue the activation of previously learned long-term memories (*Stimulation Protocol*). Various previous models with identical architecture have demonstrated how attractors can be learned via plastic BCPNN synapses (Tully et al. 2016; Lansner et al. 2013; Tully et al. 2014; Fiebig & Lansner 2017). We load each LTM network with nine orthogonal attractors (ten in the example of Figure 4B, which features two sets of five memories each). Each memory pattern consists of 16 active MCs, distributed across the 16 HCs of the network. We load-in BCPNN weights from a previously trained network (Table 2), but thereafter set *κ* = 0 to deactivate plasticity of recurrent connections in LTM stores.

In summary, the two LTM models thus feature a total of 7.46 million connections between 8.640 pyramidal cells, as well as 435.456 static connections to and from 1152 basket cells.

### Inter-area Connectivity

In our model, as in previous work, we focus on layers 2/3, as their high degree of recurrent connectivity (Thomson 2002; Yoshimura & Callaway 2005) supports attractor function. The high fine-scale specificity of dense stellate cell (Yoshimura et al. 2005) and double-bouquet cell inputs (DeFelipe et al. 2006; Chrysanthidis et al. 2018) enable strongly coding sub-populations in the superior layers of functional columns. This fits with the general observation that layers 2/3 are more input selective than the lower layers (Sakata & Harris 2009; Crochet & Petersen 2009) and thus of more immediate concern to our computational model.

The recent characterization of supragranular feedforward and feedback projections (from large cells in layer 3B and 3A, respectively), between association cortices and at short and medium cortical distances (Markov et al. 2014), allows for the construction of a basic cortical hierarchy without explicit representation of infragranular layers (and its long-range feedback projections from large cells in layer 5 and 6). This is not to say that nothing would be gained by explicitly modeling infra-granular layers, but it would go beyond the scope of this model.

Accordingly, our model implements supragranular feedforward and feedback pathways between cortical areas that are at a medium distance in the cortical hierarchy. The approximate cortical distance between Inferior Temporal Cortex (ITC) and dlPFC in macaque is about 40 mm and with an axonal conductance speed of 2 m/s, distributed conduction delays in our model (Equation 10) average just above 20 ms between these areas (Girard et al. 2001; Thorpe & Fabre-Thorpe 2001; Caminiti et al. 2013).

In the forward path, layer 3B cells in LTM project towards STM (Figure 2). We do not draw these connections one-by-one, but as branching axons targeting 25% of the pyramidal cells in a randomly chosen MC (the chance of any layer 3B cell to target any MC in STM is only 0.15%). The resulting split between targets in layer 2/3 and 4 is typical for feedforward connections at medium distances in the cortical hierarchy (Markov et al. 2014) and has important functional implications for the model (*LTM-to-STM Forward Dynamics*). We also branch off some inhibitory corticocortical connections as follows: for every excitatory connection within the selected targeted MC, an inhibitory connection is created from the same pyramidal layer 3B source cell onto a randomly selected cell outside the targeted MC, but inside the local HC. This way of drawing random forward-projections retains a degree of functional specificity due to its spatial clustering and yields patchy sparse forward-projections as observed in the cortex (Houzel et al. 1994; Voges et al. 2010) with a resulting inter-area connection probability of only 0.0125% (648 axonal projections from L3B cells to STM layers 2/3 and 4 results in ~20k total connections after branching as described above.

In the feedback path, we draw sparse plastic connections from layer 3A cells in STM to layer 2/3 cells in LTM: branching axons target 25% of the pyramidal cells in a randomly chosen HC in LTM, simulating a degree of axonal branching found in the literature (Zufferey et al. 1999). Using this method, we obtain biologically plausible sparse and structured feedback projections with an inter-area connection probability of 0.66%, which – unlike the forward pathway – do not have any built-in MC-specificity but may develop such through activity-dependent plasticity. More parameters on corticocortical projections can be found in Table 3. On average, each LTM pyramidal cell receives about 120 corticocortical connections from STM. Because about 5% of STM cells fire together during memory reactivation (see *Results*), this means that a mere 6 active synapses per target cell are sufficient for driving (and thus maintaining) LTM activity from STM (there are 96 active synapses from coactive pyramidal cells in LTM).

Notably LTMa and LTMb have no direct pathways connecting them in our model since we assume the use of previously not associated stimuli in our simulated multi-modal tasks and further, that plasticity of biological connections between them are likely too slow (LTP timescale) to make a difference in WM dynamics. This arrangement also guarantees that any binding of long-term memories across LTM areas must be the result of interaction via STM instead. Overall in our model, corticocortical connectivity is very sparse, below 1% on a cell-to-cell basis.

### Stimulation Protocol

The term *I*_*ext*_ in Equation 1 subsumes specific and unspecific external inputs. To simulate unspecific input from non-simulated columns, and other areas, pyramidal cells are continually stimulated with a zero mean noise background throughout the simulation. In each layer, two independent Poisson sources generate spikes at rate 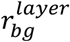, and connect onto all pyramidal neurons in that layer, via non-depressing conductances 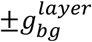 (Table 2). Before each simulation, we distribute the initial values of all plastic weights in the process of learning induced by 1.5 s low background activity (Table 2, 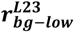). To cue the activation of a specific memory pattern (i.e. attractor), we excite LTM pyramidal cells belonging to a memory patterns component MC with an additional excitatory Poisson spike train (rate *r*_*cue*_, length *t*_*cue*_, conductance *g*_*cue*_). As LTM patterns are strongly encoded in each LTM, a brief 50 ms stimulus is usually sufficient to activate any given memory.

### Spike Train Analysis and Memory Activity Tracking

We track memory activity in time by analyzing the population firing rate of pattern-specific and network-wide spiking activity usually using an exponential moving average filter time-constant of 20 ms. We do not use an otherwise common low-pass filter with symmetrical window, because we are particularly interested in characterizing activation onsets and onset delays. As activations are characterized by sizable gamma-like bursts, a simple threshold detector can extract candidate activation events and decode the activated memory. This is trivial in LTM due to the known nature of its patterns. In STM we decode the stimulus-specificity of each cell individually by finding the maximum correlation between input pattern and the untrained STM spiking response in the 320 ms (which is the stimulation interval during plasticity-modulated stimulation period, shown in e.g. Figure 3D) following the pattern cue to LTM. Thereafter we can filter the population response of cells in STM with the same selectivity on that basis to obtain a more robust readout. We validate the specificity by means of cross-correlations, which reveal that the pattern specific populations are rather orthogonal according to the covariance matrix (off-diagonal magnitude < 0.1). In all three networks, we measure onset and offset of pattern activity by thresholding each individual activation at half of its population peak firing rate. In LTM, we further check pattern completion by analyzing component MC activation. Whenever targeted stimuli are used, we analyze peri-stimulus activation traces. When activation onsets are less predictable, such as during free STM-paced maintenance, we extract activation candidates via a threshold detector trained at the 50^th^ percentile of the cumulative distribution of the population firing rate signal.

### Synthetic field potentials and spectral analysis

We estimate local field potentials (LFPs) by calculating a temporal derivative of the average low-pass filtered (with the cut-off frequency at 250 Hz) potential for all pyramidal cells in local populations at every time step, similarly to the approach adopted by (Ursino & La Cara 2006). Although LFP is more directly linked to the synaptic activity (Logothetis 2003), the averaged membrane potentials have been reported to be correlated with LFPs (Okun et al. 2010). In particular, low-pass-filtered components of synaptic currents reflected in differentiated membrane potentials appear to carry the portion of the power spectral content of extracellular potentials that is relevant to our key findings (Lindén et al. 2010). As regards the phase response of estimated extracellular potentials, the delays of different frequency components are spatially dependent (Lindén et al. 2010). However, irrespective of the LFP synthesis, phase-related phenomena reported in this study remain qualitatively unaffected since they hinge on relative rather than absolute phase values.

Most spectral analyses have been conducted on the synthesized field potentials with the exception of population firing rates, shown in Fig. 3B and S1A. Spectral information is extracted with a multi-taper approach using a family of orthogonal tapers produced by Slepian functions (Slepian 1978; Thomson 1982), with frequency-dependent window lengths corresponding to 5-8 oscillatory cycles and frequency smoothing corresponding to 0.3-0.4 of the central frequency, which was sampled with the resolution of 1 Hz (this configuration implies that 2-3 tapers are usually used). To obtain the spectral density, spectro-temporal content is averaged within a specific time interval.

The coherence for a pair of synthesized field potentials at the spatial resolution corresponding to a hypercolumn was calculated using the multi-taper auto-spectral and cross-spectral estimates. The complex value of coherence (Carter 1987) was evaluated first based on the spectral components averaged within 0.5 s windows. Next, its magnitude was extracted to produce the time-windowed estimate of the coherence amplitude. In addition, phase locking statistics were estimated to examine synchrony without the interference of amplitude correlations (Lachaux et al. 1999; Palva 2005). In particular, phase locking value (PLV) between two signals with instantaneous phases *ϕ*_1_(*t*) and *ϕ*_2_(*t*) was evaluated within a time window of size *N*=0.5 s as follows:

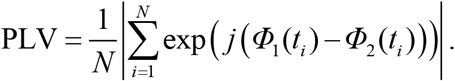

The instantaneous phase of the signals was estimated from their analytic signal representation obtained using a Hilbert transform. Before the transform was applied the signals were narrow-band filtered with low time-domain spread finite-impulse response filters (in the forward and reverse directions to avoid any phase distortions). The analysis was performed mainly for gamma-range oscillations. Continuous PLV estimate was obtained with a sliding window approach, and the average along with standard error were calculated typically over 25 trials.

### Simulation Environment and Code Accessibility

We use the NEST simulator (Gewaltig & Diesmann 2007) version 2.2 for our simulations, running on a Cray XC-40 Supercomputer of the PDC Centre for High Performance Computing. The custom-build spiking neural network implementation of the BCPNN learning rule for MPI-parallelized NEST is available on github: https://github.com/Florian-Fiebig/BCPNN-for-NEST222-MPI

### Parameter Tables

**Table 1.** Neurons, synapses, and plasticity.

|  |  |  |  |  |  |  |  |  |
| --- | --- | --- | --- | --- | --- | --- | --- | --- |
| Adaptation current | $b$ | 86 pA | Depression time constant | $\tau_{rec}$ | 500 ms | BCPNN AMPA gain | $w_{gain}^{AMPA}$ | 3.93 nS |
| Adaptation time constant | $\tau_{Iw}$ | 500 ms | AMPA synaptic time constant | $\tau^{AMPA}$ | 5 ms | BCPNN NMDA gain | $w_{gain}^{NMDA}$ | 0.21 nS |
| Membrane Capacity | $C_m$ | 280 pF | NMDA synaptic time constant | $\tau^{NMDA}$ | 100 ms | BCPNN bias current gain | $\beta_{gain}$ | 90 pA |
| Leak Reversal Potential | $E_L$ | -70 mV | GABA synaptic time constant | $\tau^{GABA}$ | 5 ms | BCPNN lowest rate | $f_{min}$ | 0.2 Hz |
| Leak Conductance | $g_L$ | 14 pS | AMPA Reversal Potential | $E^{AMPA}$ | 0 mV | BCPNN highest rate | $f_{max}$ | 20 Hz |
| Upstroke slope factor | $\Delta_T$ | 3 mV | NMDA Reversal Potential | $E^{NMDA}$ | 0 mV | BCPNN lowest probability | $\epsilon$ | 0.01 |
| Spike Threshold | $V_t$ | -55 mV | GABA Reversal Potential | $E^{GABA}$ | -75 mV | BCPNN Spike event duration | $\Delta t$ | 1 ms |
| Spike Reset Potential | $V_r$ | -80 mV | Dopaminergic Modulation | $\kappa_p$ | 6.0 | P-Trace time constant | $\tau_p$ | 5 s |
| Utilization factor | $U$ | .33 | Regular Plasticity | $\kappa_{normal}$ | 1.0 | | | |

**Table 2.** Network size, Conduction delay, Stimulation, LTM Preload BCPNN weights. Layer 4 not simulated in LTM.

|  |  |  |  |  |  |  |
| --- | --- | --- | --- | --- | --- | --- |
| STM patch size | 17 x 17 mm | | Initialization Input rate layer 2/3 | | $r_{bg-low}^{L23}$ | 550 Hz |
| Simulated HCs | $n_{HC}^{STM}$ | 25 | Background activity rate layer 2/3 | | $r_{bg}^{L23}$ | 625 Hz |
| Simulated MC per HC | $n_{MC}^{STM}$ | 12 | Background activity rate layer 4 | | $r_{bg}^{L4}$ | 300 Hz |
| LTM patch size | 25 x 25 mm | | High Background activity rate layer 2/3 (e.g. STM Maintainance) | | $r_{bg-high}^{L23}$ | 950 Hz |
| Simulated HCs | $n_{HC}^{LTM}$ | 16 | | | | |
| Simulated MC per HC | $n_{MC}^{LTM}$ | 9 | Background conductance | | $g_{bg}$ | $\pm 1.5$ nS |
| Axonal Conduction Speed | $V$ | $2 \frac{m}{s}$ | | | | |
| Minimal conduction delay | $t_{min}^{syn}$ | 1.5 ms | Cue stimulus duration | | $t_{cue}$ | 50 ms |
| STM – LTM distance | $d_{STM-LTM}$ | 40 mm | Stimulation rate | | $r_{cue}$ | 650 Hz |
| Hypercolumn diameter | $d_{HC}$ | 0.64 mm | Cue stimulus conductance | | $g_{cue}$ | +1.5 nS |
| Layer 2 pyramidal per MC | $n_{MC}^{PYR-L2}$ | 20 | LTM Intra HC – Intra MC weight | | $w_{IntraMC}^{IntraHC}$ | $3.36 w_{gain}^{syn}$ |
| Layer 3A pyramidal per MC | $n_{MC}^{PYR-L3A}$ | 5 | LTM Intra HC – Inter MC weight | | $w_{InterMC}^{IntraHC}$ | $-4.82 w_{gain}^{syn}$ |
| Layer 3B pyramidal per MC | $n_{MC}^{PYR-L3B}$ | 5 | | | | |
| Layer 4 pyramidal per MC | $n_{MC}^{PYR-L4}$ | 30 | LTM Inter HC – Coactive MC weight | | $w_{CoactiveMC}^{InterHC}$ | $3.08 w_{gain}^{syn}$ |
| Basket cells per MC | $n_{MC}^{basket}$ | 4 | LTM Inter HC – Competing MC weight | | $w_{CompetingMC}^{InterHC}$ | $-4.28 w_{gain}^{syn}$ |

**Table 3.** Projections.

| Scope | Source | Target | Type | Symbol | Value |
| --- | --- | --- | --- | --- | --- |
| Cortical Area | Pyramidal | Basket | probability | $p_{P-B}$ | 0.7 |
| | Pyramidal | Basket | conductance (static) | $g_{P-B}$ | +3.5 nS |
| | Basket | Pyramidal | probability | $p_{B-P}$ | 0.7 |
| | Basket | Pyramidal | conductance (static) | $g_{B-P}$ | -20 nS |
| | L23e | L23e | probability | $p_{L23e-L23e}$ | 0.2 |
| | L23e | L23e | AMPA gain (BCPNN) | $w_{gain}^{AMPA}$ | 3.93nS |
| | L23e | L23e | NMDA gain (BCPNN) | $w_{gain}^{NMDA}$ | 0.21nS |
| | L4e | L23e | probability | $p_{L4e-L23e}$ | 0.25 |
| | L4e | L23e | conductance (static) | $g_{L4e-L23e}$ | 25 nS |
| Feed forward | LTM L3Ae | STM MC | probability | $p_{L3Ae-MC}^{FF}$ | 0.0015 |
| | LTM L3Ae | STM MC | branching factor | $b_{L3Ae-MC}^{FF}$ | 0.25 |
| | LTM L3Ae | STM L23e | conductance (static) | $g_{L3Ae-L23e}^{FF}$ | $\pm 7.2$ nS |
| | LTM L3Ae | STM L4e | conductance (static) | $g_{L3Ae-L4e}^{FF}$ | $\pm 7.2$ nS |
| Feedback | STM PYR | LTM PYR | probability | $p_{P-P}^{FB}$ | 0.0066 |
| | STM L3Be | LTM HC | branching factor | $b_{L3Be-HC}^{FB}$ | 0.25 |
| | STM L3Be | LTM L23e | AMPA gain (BCPNN) | $w_{FB}^{AMPA}$ | 7.07 nS |
| | STM L3Be | LTM L23e | NMDA gain (BCPNN) | $w_{FB}^{NMDA}$ | 0.4 nS |

## Results

Our model implements WM function arising from the interaction of STM and LTM networks, which manifests itself in multi-modal memory binding phenomena. To this end, we simulate three cortical patches with significant biophysical detail: one STM and two LTM networks (LTMa, LTMb), representing PFC and parieto-temporal areas, respectively (Figure 2). The computational network model used here represents a detailed modular cortical microcircuit architecture in line with previous models (Lundqvist, Rehn, Djurfeldt, & Lansner, 2006; Tully et al., 2016; Lundqvist et al., 2011). Like those models, the new model can reproduce a wide range of meso- and macroscopic biological manifestations of cortical memory function including complex oscillatory dynamics and synchronization effects (Lundqvist et al. 2011; Lundqvist et al. 2013; Silverstein & Lansner 2011). The current model is built directly upon a recent STM model of human word-list learning (Fiebig & Lansner 2017). The associative cortical layer 2/3 network of that model was sub-divided into layers 2, 3A, and 3B. Importantly, in this work we extend this model with an input layer 4 and corticocortical connectivity to LTM stores in temporal cortical regions. This large, multi-area network model synthesizes many different anatomical and electrophysiological cortical data and produces complex output dynamics. Here, we specifically focus on the dynamics of memory specific subpopulations in the interaction of STM and LTM networks.

We introduce the operation of the WM model in several steps. First, we take a brief look at background activity and active memory states in isolated cortical networks of this kind to familiarize the reader with some of its dynamical properties. Second, we describe the effect of memory activation on STM with and without plasticity. Third, we add the plastic backprojections from STM to LTM and monitor the encoding and maintenance of several memories in the resulting STM-LTM loop. We track the evolution of acquired cell assemblies with shared pattern-selectivity in STM and show their important role in WM maintenance (aka delay activity). We then demonstrate that the emerging WM network system is capable of flexibly updating the set of maintained memories. Finally, we simulate multi-modal association and analyze its dynamical correlates. We explore temporal characteristics of network activations, the accompanying oscillatory behavior of the synthesized field potentials, cross-cortical delays as well as gamma-band coupling (coherence and phase synchronization) between LTM networks during WM encoding, maintenance, and cue-driven associative recall of multi-modal memories (LTMa-LTMb pairs of associated memories).

**Figure 3.**
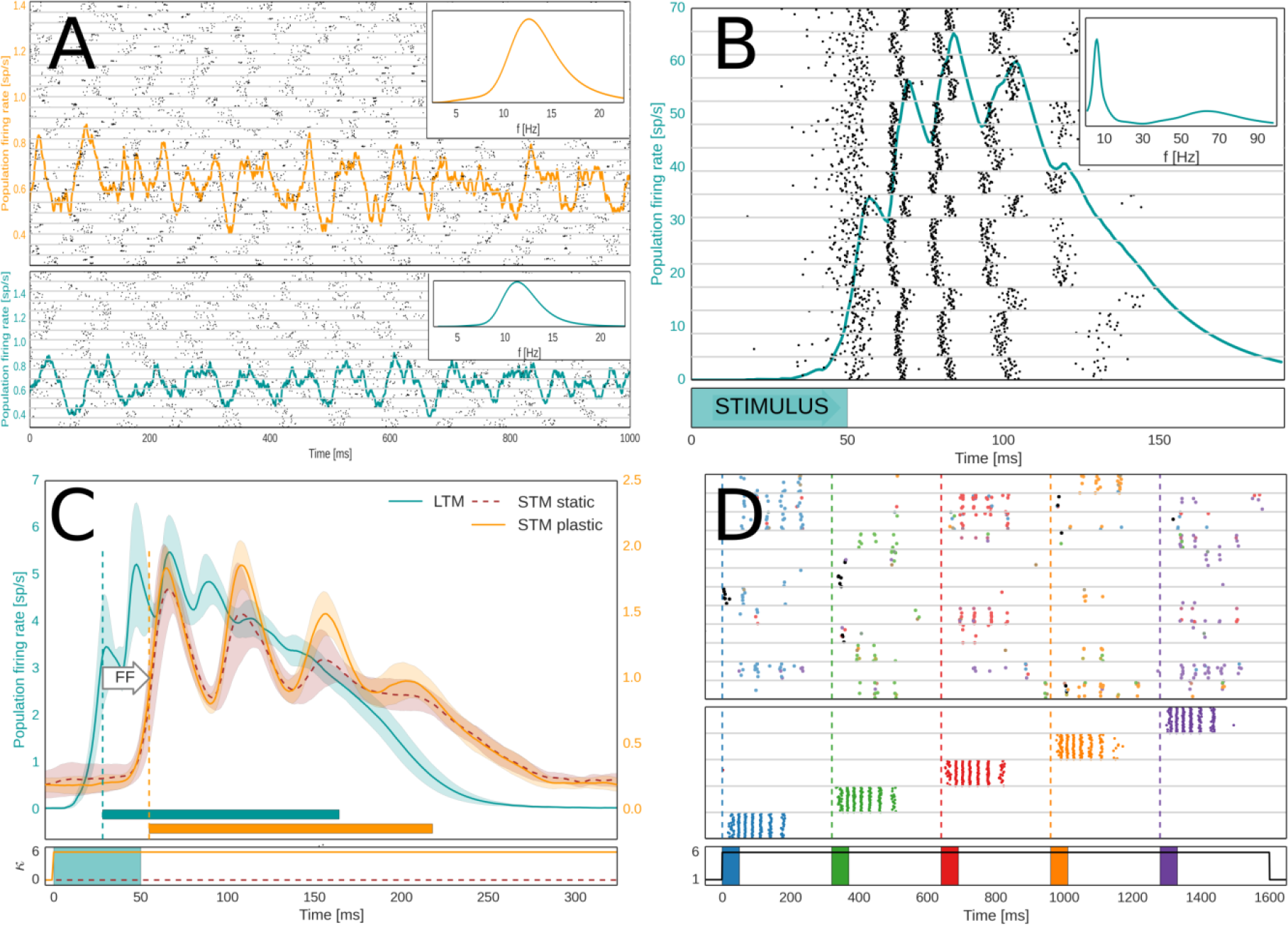
Basic Network behavior in spike rasters and population firing rates. A: Activity in the untrained network under strong background input. **A:** Subsampled spike raster of STM (top) and LTM (bottom) layer 2/3 activity. HCs are separated by grey horizontal lines. Global oscillations in the alpha range (10-13 Hz) characterize this activity state in both STM (top) and LTM (bottom) in the absence of attractors. Inset: Power Spectral Density of each network’s LFP. **B:** Cued LTM memory activation express as fast oscillation bursts of selective cells (50-80 Hz), organized into a theta-like envelope (4-8 Hz), see also Power Spectrum Inset. The gamma-band is broad due to varying length of the underlying cycles, i.e. noticeably increasing over the short memory activation period. The underlying spike raster shows layer 2/3 activity of the activated MC in each HC, revealing spatial synchronization. The brief stimulus is a memory specific cue. **C:** LTM-to-STM forward dynamics as shown in population firing rates of STM and LTM activity following LTM-activation induced by a 50 ms targeted stimulus at time 0. LTM-driven activations of STM are characterized by a feedforward delay (FF). Shadows indicate the standard deviation of 100 peri-stimulus activations in LTM (blue) and STM with (orange) and without plasticity enabled (dashed, dark orange). Horizontal bars indicate the activation half-width (*Methods*). Onset is denoted by vertical dashed lines. The stimulation of LTM and activation of plasticity is denoted underneath. **D:** Subsampled spike raster of STM (top) and LTM (middle) during forward activation of the untrained STM by five different LTM memory patterns, triggered via specific memory cues in LTM at times marked by the vertical dashed lines. Bottom spike raster shows LTM layer 2/3 activity of one selective MC per activated pattern (colors indicate different patterns). Top spike raster shows layer 2/3 activity of one HC in STM. STM spikes are colored according to each cells dominant pattern-selectivity (based on the memory pattern correlation of individual STM cell spiking during initial pattern activation, see Methods, *Spike Train Analysis and Memory Activity Tracking*). Bottom: The five stimuli to LTM (colored boxes) and modulation of STM plasticity (black line).

### Background activity and Activated memory

The untrained network (see *Methods*) features fluctuations in membrane voltages and low-rate, asynchronous spiking activity (Figure 3 – Supplement 1). At higher background input levels, the empty network transitions into a state characterized by global oscillations of the population firing rates in the alpha/beta range (Figure 3A). This is largely an effect of fast feedback inhibition from local basket cells (Figure 1), high connection density within MCs, and low latency local spike transmission (Lundqvist et al. 2010). If the network has been trained with structured input so as to encode memory (i.e. attractor states), background noise or a specific cue (*Methods*) can trigger memory item reactivations accompanied by fast broad-band oscillations modulated by an underlying slow oscillation in the lower theta range (~4-8 Hz) (Lundqvist et al. 2011; Herman et al. 2013) (Figure 3B). The spiking activity of memory activations (aka attractors) is short-lived due to neural adaptation and synaptic depression. When unspecific background excitation is very strong, this can result in a random walk across stored memories (Fiebig & Lansner 2017; Lundqvist et al. 2011).

### LTM-to-STM Forward Dynamics

We now consider cued activation of several memories embedded in LTM. Each HC in LTM features selectively coding MCs for given memory patterns that activate synchronously in theta-like cycles each containing several fast oscillation bursts (Figure 3B). Five different LTM memory patterns are triggered by brief cues, accompanied by an upregulation of STM plasticity, see Figure 3D **(bottom)**. To indicate the spatio-temporal structure of evoked activations in STM, we also show a simultaneous subsampled STM spike raster (Figure 3D **top**). STM activations are sparse (ca 5%), but despite this, nearby cells (in the same MC) often fire together. The distributed, patchy character of the STM response to memory activations (Figure 3D **top**) is shaped by branching forward-projections from LTM layer 3B cells, which tend to activate close-by cells. STM input layer 4 receives half of these corticocortical connections and features very high fine-scale specificity in its projections to layer 2/3 pyramidal neurons, which furthers recruitment of local clusters with shared selectivity. STM cells initially fire less than those in LTM because the latter received a brief, but strong activation cue and have strong recurrent connections if they code for the same embedded memory pattern. STM spikes in Figure 3D are colored according to the cells’ dominant memory pattern selectivity (Methods, *Spike Train Analysis and Memory Activity Tracking*), which reveals that STM activations are mostly non-overlapping as well. Unlike the organization of LTM with strictly non-overlapping memory patterns, MC activity in STM is not exclusive to any given input pattern. Nevertheless, nearby STM cells often develop similar pattern selectivity. On the other hand, different stimulus patterns typically develop quite non-overlapping STM representations. This is due to the randomness in feed-forward LTM to STM connectivity, competition via basket cell feedback inhibition, and short-term dynamics, such as neural adaptation and synaptic depression. STM neurons that have recently been activated by a strong, bursting input from LTM are refractory and thus less prone to spike again for some time thereafter (*τ*_*rec*_ and 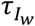, Table 1), further reducing the likelihood of creating overlapping STM representations for different patterns.

Figure 3C shows peri-stimulus population firing rates of both STM and LTM networks (mean across 100 trials with five triggered memories each). There is a bottom-up response delay between stimulus onset at t=0 and LTM activation, as well as a substantial forward delay. Oscillatory activity in STM is lower than in LTM mostly because the untrained STM lacks strong recurrent connections. It is thus less excitable, and therefore does not trigger its basket cells (the main drivers of fast oscillations in our model) as quickly as in LTM. Fast oscillations in STM and the amplitude of their theta-like envelope build up within a few seconds as new cell assemblies become stronger (e.g. Figure 4A and Figure 4 - Supplement 1). As seen in Figure 3B, bursts of co-activated MCs in LTM can become asynchronous during activation. Dispersed forward axonal conduction delays further decorrelate this gamma-like input to STM. Activating strong plasticity in STM (*κ* = *κ*_*p*_, *Methods* and Table 1) has a noticeable effect on the amplitude of stimulus-locked oscillatory STM activity after as little as 100 ms (cf. Figure 3C, **STM**).

**Figure 4.**
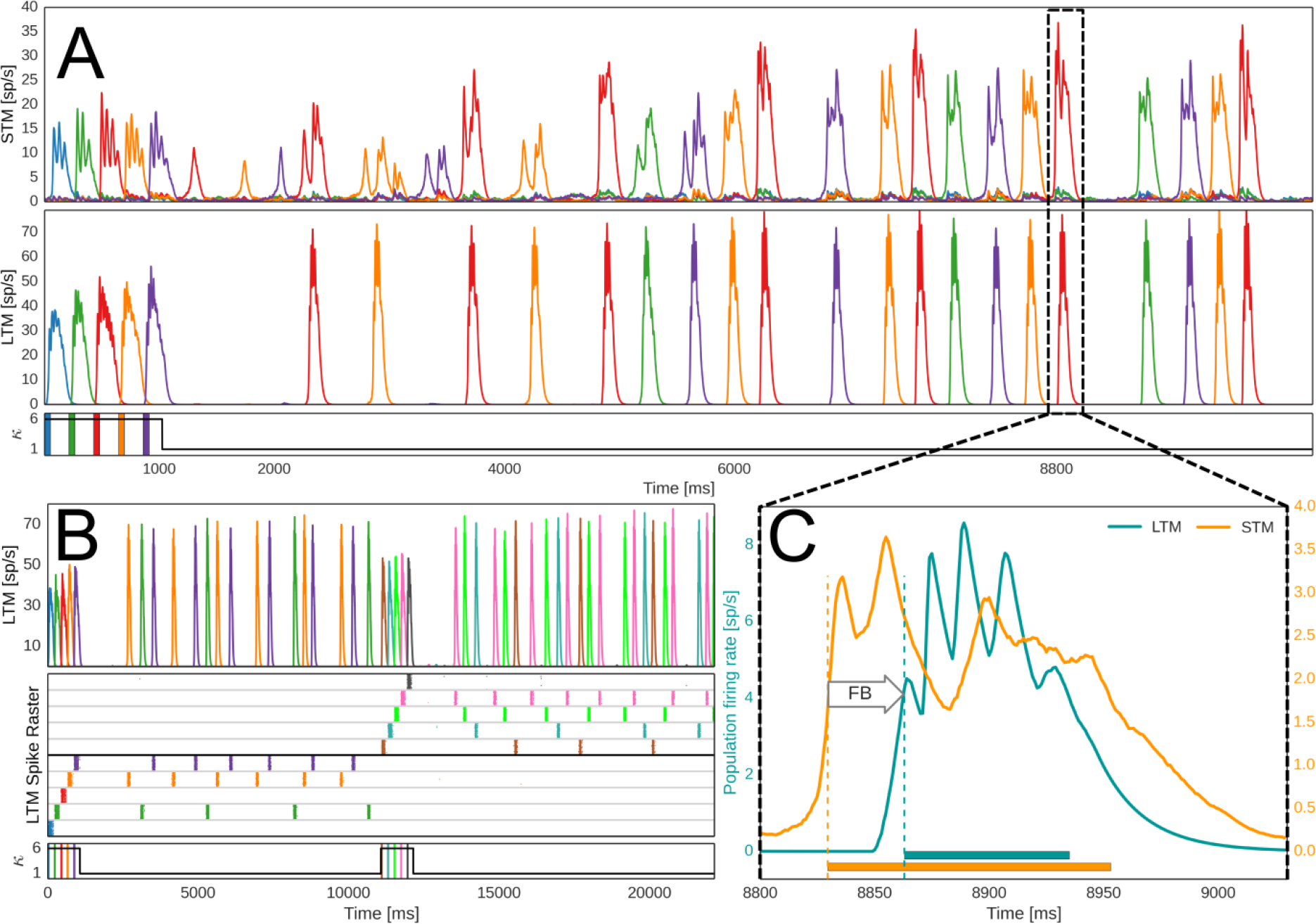
Encoding and feedback-driven reactivation of LTM. **A**: Firing rates of pattern-specific subpopulations in STM and LTM during encoding and subsequent maintenance of five memories. Just as in the plasticity-modulated stimulation phase shown in Figure 2D, five LTM memories are cued via targeted 50 ms stimuli (shown underneath). Plasticity of STM and its backprojections is again elevated six-fold during the initial memory activation. Thereafter, a strong noise drive to STM causes spontaneous activations and plasticity induced consolidation of pattern-specific subpopulations in STM (lower plasticity, *κ* = 1). Backprojections from STM cell assemblies help reactivate associated LTM memories. **B:** Updating of WM. Rapid encoding and subsequent maintenance of a second group of memories following an earlier set. The LTM spike raster shows layer 2/3 activity of one LTM HC (MCs separated by grey horizontal lines), the population firing rate of pattern-specific subpopulations across the whole LTM network is seen above. Underneath we denote stimuli to LTM and the modulation of plasticity, *κ*, in STM and its backprojections. **C:**STM-to-LTM loop dynamics during a spontaneous reactivation event. STM-triggered activations of LTM memories are characterized by a feedback delay and a second peak in STM after LTM activations. Horizontal bars at the bottom indicate activation half-width (*Methods*). Onset is denoted by vertical dashed lines.

### Multi-item Working Memory

In Figure 3D, we have shown pattern-specific subpopulations in STM emerging from feedforward input. Modulated STM plasticity allows for the quick formation of rather weak STM cell assemblies from one-shot learning. When we include plastic STM backprojections, these assemblies can serve as an index for specific LTM memories and provide top-down control signals for memory maintenance and retrieval. STM backprojections with fast Hebbian plasticity can index multiple activated memories in the closed STM-LTM loop. In Figure 4A, we show network activity following targeted activation of five LTM memories (Spike raster in Figure 4 - Supplement 1). Under an increased unspecific noise-drive (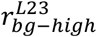, Table 2), STM cell assemblies, formed during the brief plasticity-modulated stimulus phase (cf. Figure 3D) may activate spontaneously. These brief bursts of activity are initially weak and different from the theta-like cycles of repeated fast bursting seen in LTM attractor activity.

STM recurrent connections remain plastic (*κ* = 1) throughout the simulation, so each reactivation event further strengthens memory-specific cell assemblies in STM. As a result, there is a noticeable ramp-up in the strength of STM pattern-specific activity over the course of the delay period (cf. increasing burst length and amplitude in Figure 4A, or Figure 4 - Supplement 2). STM backprojections are also plastic and thus acquire memory specificity from STM-LTM co-activations, especially during the initial stimulation phase. Given enough STM cell assembly firing, their sparse but potentiated backprojections can trigger associated memories in LTM. Weakly active assemblies may fail to do so. In the example of Figure 4A, we can see a few early STM reactivations that are not accompanied (or quickly followed) by a corresponding LTM pattern activation (of the same color) in the first two seconds after the plasticity-modulated stimulation phase. When LTM is triggered, there is a noticeable feedback delay (Figure 4C), which we will address together with aforementioned feed forward delays in the analysis of recall dynamics during a multi-item, multi-modal recall task.

Cortical feedforward and feedback pathways between LTM and STM form a loop, so each LTM activation will again feed into STM, typically causing a second peak of activation in STM 40 ms after the first (Figure 4C). The forward delay from LTM to STM, which we have seen earlier in the stimulus-driven input phase (Figure 3C), is still evident here in this delayed secondary increase of the STM activation following LTM onset. The reverberating cross-cortical activation extends/sustains the memory activation and thus helps stabilize item-specific STM cell assemblies and their specificity. This effect may be called auto-consolidation and it is an emergent feature of the plastic STM-LTM loop in our model. It occurs on a timescale governed by the unmodulated plasticity time constant (*κ* = *κ*_*normal*_, *τ*_*p*_ = 5 *s*, Table 1). After a few seconds, the network has effectively stabilized and typically maintains a small set of 3-4 activated long-term memories. The closed STM-LTM loop thus constitutes a functional multi-item WM.

A crucial feature of any WM system is its flexibility, and Figure 3B highlights an example of rapid updating. The maintained set of activated memories can be weakened by stimulating yet another set of input memories. Generally speaking, earlier items are reliably displaced from active maintenance in our model if activation of the new items is accompanied by the same transient elevation of plasticity (*κ*_*p*_/*κ*_*normal*_, Table 1) used during the original encoding of the first five memories (Corresponding population firing rates and spike rasters are shown in Figures 4 - Supplements 2,3).

In line with the earlier results (Fiebig & Lansner 2017), cued activation can usually still retrieve previously maintained items. The rate of decay for memories outside the maintained set depends critically on the amount of noise in the system, which erodes the learned associations between STM and LTM neurons as well as STM cell assemblies. We note that such activity-dependent memory decay is substantially different from time-dependent decay, as shown by Mi et al.(2017).

### Multi-modal, Multi-item Working Memory

Next, we explore the ability of the closed STM-LTM loop system to flexibly bind co-active pairs of long-term memories from different modalities (LTMa and LTMb, respectively). As both LTM activations trigger cells in STM via feedforward projections, a unique joint STM cell assembly with shared pattern-selectivity is created. Forward-activations include excitation and inhibition and combine non-linearly with each other (*Methods*) and with prior STM content.

Figure 5 illustrates how this new index then supports WM operations, including delay maintenance through STM-paced co-activation events and stimulus-driven associative memory pair completion. The three columns of Figure 5 illustrate three fundamental modes of the closed STM-LTM loop: stimulus-driven encoding, WM maintenance, and associative recall. The top three rows show sampled activity of a single trial (see also Figure 5 – Supplement 1), whereas the bottom row shows multi-trial averages.

**Figure 5.**
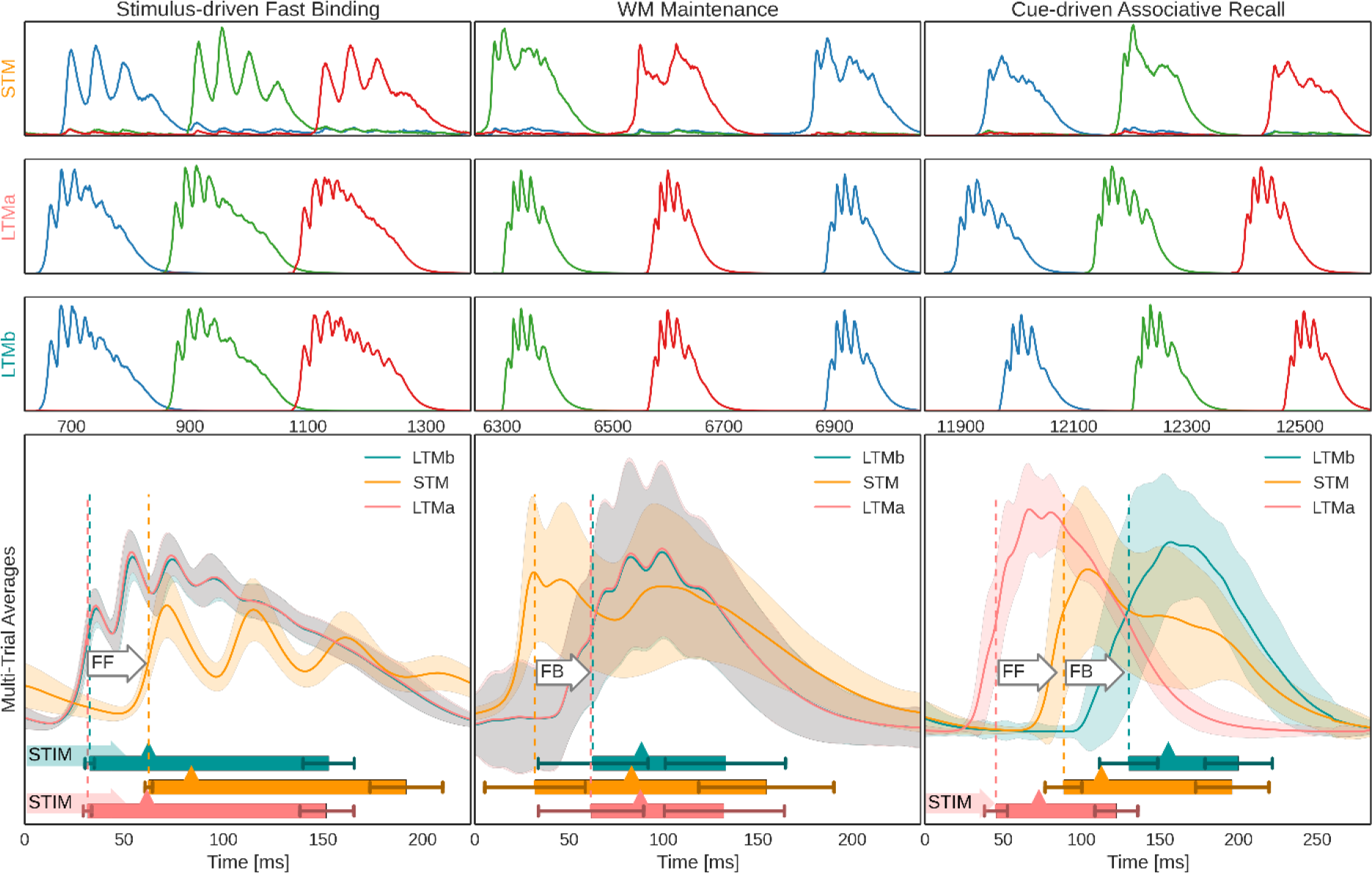
Population firing rates of networks and memory-specific subpopulations during three different modes of network activity. **Top-Half**: Exemplary activation of three memories (blue, green, red respectively) in STM (1^st^ row), LTMa (2^nd^ row), and LTMb (3^rd^ row) during three different modes of network activity: The initial association of pairs of LTM memory activations in STM (left column), WM Maintenance through spontaneous STM-paced activations of bound LTM memory pairs (middle column), and cue-driven associative recall of previously paired stimuli (right column). **Bottom-Half**: Multi-trial peri-stimulus activity traces from the three cortical patches across 100 trials (495 traces, as each trial features 5 activated and maintained LTM memory pairs and very few failures of paired activation). Shaded areas indicate a standard deviation from the underlying traces. Vertical dashed lines denote mean onset of each network’s activity, as determined by activation half-width (*Methods*), also denoted by a box underneath the traces. Error bars indicate a standard deviation from activation onset and offset. Mean peak activation is denoted by a triangle on the box, and shaded arrows to the left of the box denote targeted pattern stimulation of a network at time 0. As there are no external cues during WM maintenance (aka delay period), we use detected STM activation onset to align firing rate traces of 5168 STM-paced LTM-reactivations across trials and reactivation events for averaging. White arrows annotate feedforward (FF) and feedback (FB) delay, as defined by respective network onsets.

During stimulus-driven association, we co-activate memories from both LTMs by brief 50 ms cues that trigger activation of the corresponding memory patterns. The average of peri-stimulus activations reveals 45 ± 7.3 ms LTM attractor activation delay, followed by 43 ± 7.8 ms feedforward delay (about half of which is explained by axonal conduction delays due to the spatial distance between LTM and STM) from the onset of the LTM activations to the onset of the input-specific STM response (Figure 5 **top-left** and **bottom-left**).

During WM maintenance, a 10 s delay period, paired LTM memories reactivate together. Onset of these paired activations is a lot more variable than during cued activation with a feedback delay mean of 41.5 ± 15.3 ms, mostly because the driving STM-activations are of variable size and strength. Over the course of the maintenance period the oscillatory dynamics of the LTMs changes. In particular, LFP spectral power as well as coherence between LTMs in the broad gamma (30-80 Hz) band increases (p<0.001 for each of two permutation tests comparing average spectral power/coherence in the gamma band between two intervals during the delay period: 4-8 s and 8-12 s; n=25 trials). To study the fast oscillatory dynamics of the LFP interactions between LTMs during the WM maintenance, mediated by STM, we follow up the coherence analysis and examine the gamma phase synchronization effect using PLV with 0.5 s sliding window (see *Methods*). It appears that the gamma phase coupling also increases during the second part of the WM maintenance period (p<0.001 in analogous permutation test as above; Figure 6).

**Figure 6.**
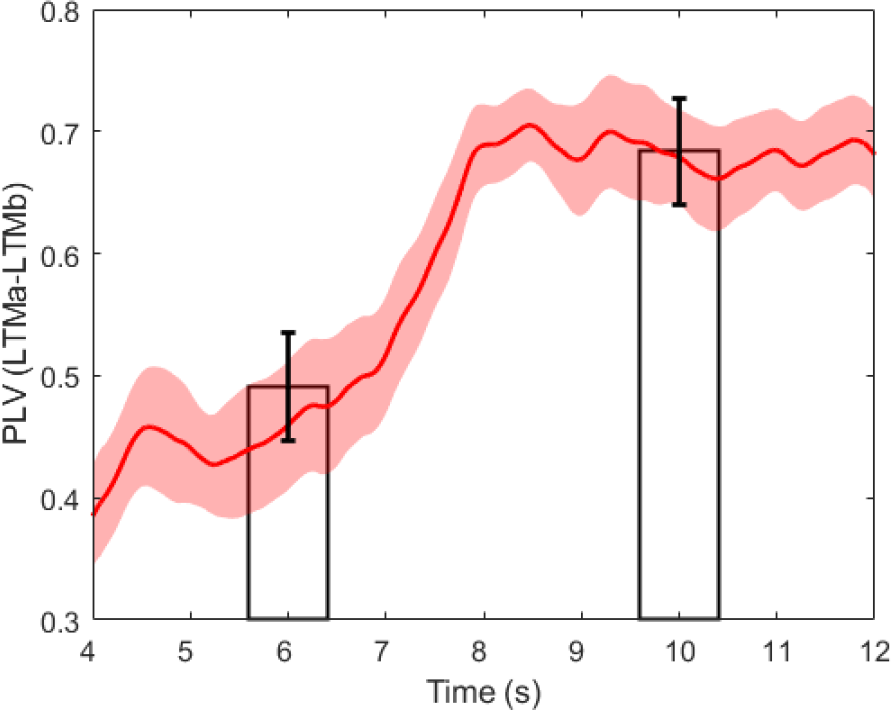
Gamma-band Phase Locking Value (PLV) between LTMa and LTMb during WM maintenance. PLV is estimated using sliding window of size 0.5 s (the period between 4 and 12 s is shown). Two bars demonstrate the average gamma-band PLV over the first (4-8 s) and the second part (8-12 s) of the WM maintenance period. Shaded area and error bars correspond to the standard error of the mean calculated over n=25 trials.

Following the maintenance period, we test the memory system’s ability for bi-modal associative recall. To this end, we cue LTMa, again using a targeted 50 ms cue for each memory, and track the systems response across the STM-LTM loop. We compute multi-trial averages of peri-stimulus activations during recall testing (Figure 5 **bottom-right**). Following cued activation of LTMa, STM responds with the related joint cell assembly activation as the input is strongly correlated to the learned inputs, as a result of the simultaneous activation with LTMb earlier on. Similar to the mnemonic function of an index, the completed STM pattern then triggers the associated memory in LTMb through backprojections. STM activation now extends far beyond the transient activity of LTMa because STM recurrent connectivity and the STM-LTMb recurrence re-excites it. Temporal overlap between associated LTMa and LTMb memory activations peaks around 125 ms after the initial stimulus to LTMa.

### Network Power Spectra and the Non Associative Control Case

**Figure 7.**
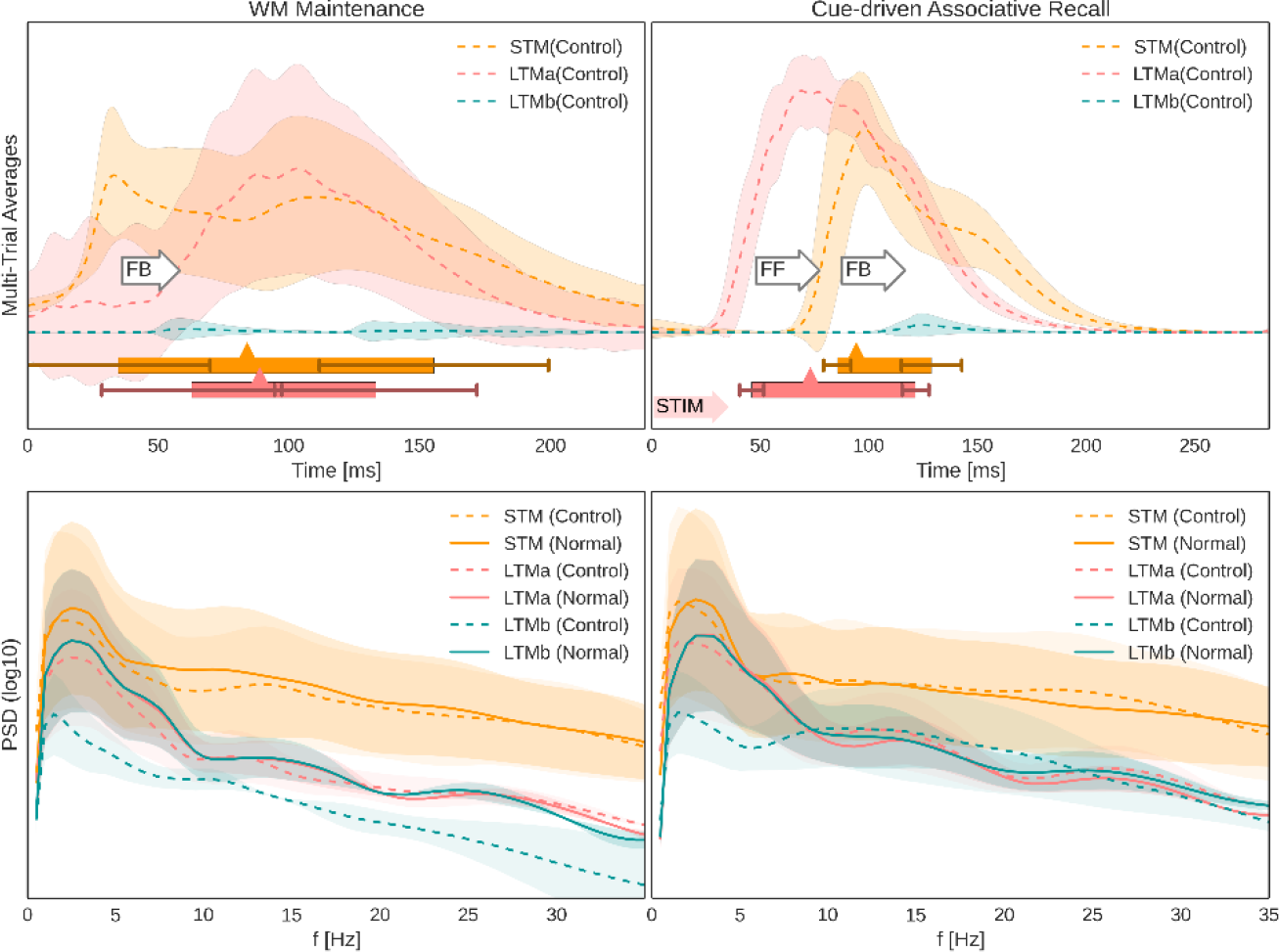
Non-Associative Control Case and Power Spectral Analysis: **Top-Half**: Multi-trial peri-stimulus activity traces from the three cortical patches across 25 trials following WM-encoded LTMa activations as before, but without associated LTMb memory activations. Shaded areas indicate a standard deviation from the underlying traces. Activation half-widths (*Methods*) denoted by a box underneath the traces. Error bars indicate a standard deviation from activation onset and offset. Mean peak activation is denoted by a triangle on the box, and shaded arrows to the left of the box denote targeted pattern stimulation of LTMa at time 0. As there are no external cues during WM maintenance (aka delay period), we use detected STM activation onset to align firing rate traces of 406 STM-paced LTMa-reactivations across trials and reactivation events for averaging. There is no evidence of associated LTMb activations in the control case (only small increases in spike rate variability). White arrows annotate feedforward (FF) and feedback (FB) delay, as defined by respective network onsets. **Bottom-Half**: Power spectral density of synthesized LFPs estimated over the maintenance (left) and recall (right) periods for STM and both LTMs in two cases: with (solid lines) and without (dashed line; control case) associated LTMb memory activations. Please note the log-scale. Shaded areas correspond to the standard deviation of the mean PSD over 25 trials. The decrease in theta- and gamma-band power observed during the maintenance (left) and recall (right) periods in the LTMb in the control case is due to lack of memory pattern reactivations in LTMb as they are not associated with LTMa via STM.

Figure 7 **(top)** shows multi-trial peri-stimulus/peri-activation activity traces for a control task, where learned and maintained LTMa items are not associated with concurrent LTMb activations. LTMa items are still encoded in STM, maintained over the delay, and recalled by specific cues, but LTMb now remains silent throughout the maintenance period (Figure 7 **top-left**) and as expected does not show any evidence of memory activation following LTMa-specific cues during recall testing (Figure 7 **top-right**, see also LFP signal in Figure 7 – Supplement 2). The logarithmic power spectra (Figure 7 **bottom**) show a noticeable difference between the normal associative and the non-associative control trials. The latter displays a significant drop in LTMb power across the board, particularly during the maintenance period. This can be explained by the overall lower number of memory reactivations in STM during the non-associative control task (2.58±0.28 vs 1.62±0.47 reactivations/s).

### Top-Down and Bottom-Up Delays

We collected distributions of feedforward and feedback delays during associative recall (Figure 8). To facilitate a more immediate comparison with biological timing data we also computed the Bottom-Up and Top-Down response latency of the model in analogy to Tomita et al. (1999). Their study explicitly tested widely held beliefs about the executive control of PFC over ITC in memory retrieval. To this end, they identified and recorded neurons in ITC of monkeys trained to memorize several visual stimulus-stimulus associations. They employed a posterior-split brain paradigm to cleanly disassociate the timing of the bottom-up (contralateral stimuli) and top-down response (ipsilateral stimuli) in 43 neurons significantly stimulus-selective in both conditions. They observed that the latency of the top-down response (178 ms) was longer than that of the bottom-up response (73 ms).

**Figure 8.**
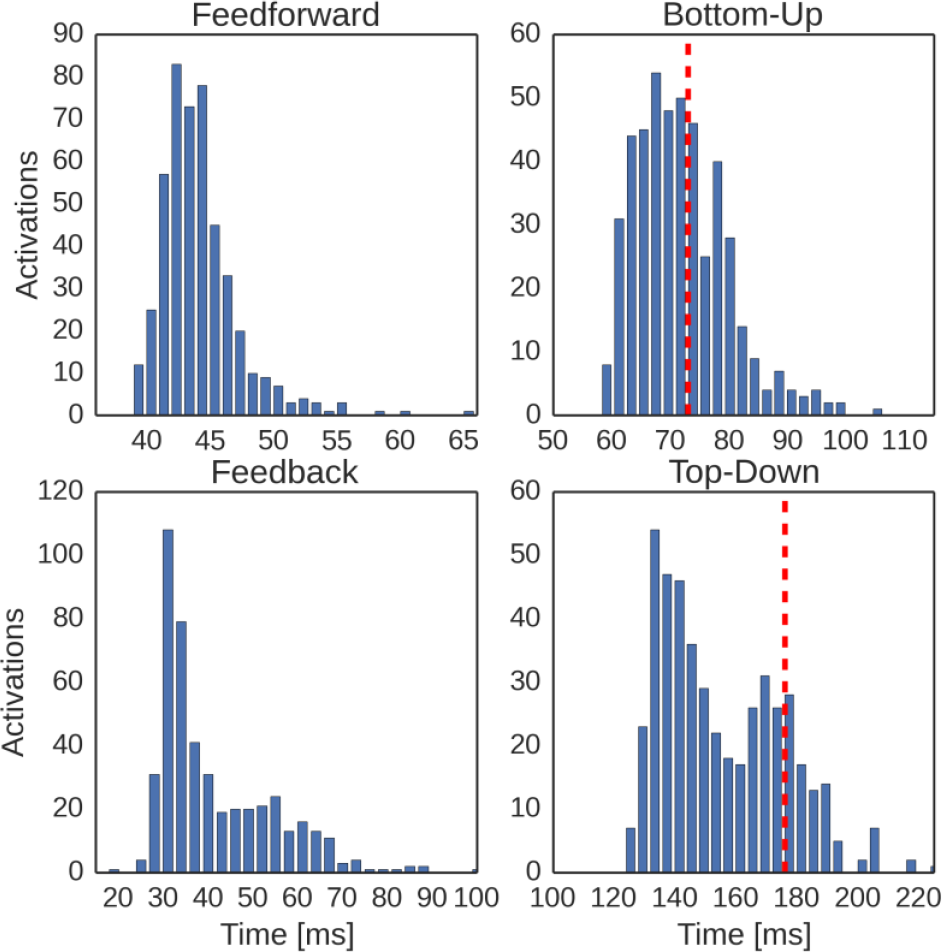
Comparison of key activation delays during associative recall in model and experiment following a cue to LTMa. **Top-Left**: Feedforward delay distribution in the model, as defined by the temporal delay between LTMa onset and STM onset (as shown in Figure 4, Bottom-right). **Top-Right**: Bottom-up delay distribution in the model, as defined by the temporal delay between stimulation onset and LTMa peak activation. The red line denotes the mean bottom-up delay, as measured by Tomita et al. (1999). **Bottom-Left**: Feedback delay distribution in the model, as defined by the temporal delay between STM onset and LTMb onset (measured by half-width, as shown in Figure 4, Bottom-right). **Bottom-Right**: Top-Down delay distribution in the model, as defined by the temporal delay between stimulation onset and LTMb peak activation. The red line denotes the mean bottom-up delay, as measured by Tomita et al. (1999). Model delays were averaged over 100 trials with 5 paired stimuli each.

Our simulation is analogous to this experimental setup with respect to some key features, such as the spatial extent of memory areas (STM/dlPFC about 289 mm^2^) and inter-area distances (40 mm cortical distance between PFC and ITC). These measures heavily influence the resulting connection delays and time needed for information integration. In analogy to the posterior-split brain experiment, our model’s LTMa and LTMb are unconnected. However, we now have to consider them as ipsi- and contralateral visual areas in ITC. The display of a cue in one hemi-field in the experiment then corresponds to the LTMa-sided stimulation of an associated memory pair in the model. This arrangement forces any LTM interaction through STM (representing PFC), and allows us to treat the cued LTMa memory activation as a Bottom-up response, whereas the much later activation of the associated LTMb representation is related to the Top-down response in the experimental study. Figure 8 shows the distribution of these latencies in our simulations, where we also marked the mean latencies measured by Tomita et al. The mean of our bottom-up delay (72.9 ms) matches the experimental data (73 ms), whereas the mean of the broader top-down latency distribution (155.2 ms) is a bit lower than in the monkey study (178 ms). Of these 155.2 ms, only 48 ms are explained by the spatial distance between networks, as verified by a fully functional alternative model with 0 mm distance between networks.

## Discussion

In this work, we have proposed and studied a novel theory for WM that rests on the dynamic interactions between STM and LTM stores shaped by fast synaptic plasticity. In particular, it hypothesizes that activity in parieto-temporal LTM stores targeting PFC via fixed or slowly plastic and patchy synaptic connections triggers an activity pattern in PFC, which then gets rapidly encoded by means of fast Hebbian plasticity to form a cell assembly. Equally plastic backprojections from PFC to the LTM stores are enhanced as well, thereby associating the formed PFC “index” specifically with the active LTM cell assemblies. This rapidly but temporarily enhanced connectivity produces a functional WM system capable of encoding and maintaining multiple individual LTM items, i.e. bringing these LTM representations “on-line”, and forming novel associations within and between several connected LTM areas and modalities. The PFC cell assemblies themselves do not encode much information but act as indices into LTM stores, which contain additional information that is also more permanent. The underlying highly plastic connectivity and thereby the WM itself is flexibly remodeled and updated as new incoming activity gradually over-writes previous WM content.

We have studied the functional and dynamical implications of this theory by implementing and evaluating a special case of a biologically plausible large-scale spiking neural network model representing PFC reciprocally connected with two LTM areas (visual and auditory) in temporal cortex. We have shown how a number of single LTM items can be encoded and maintained “on-line” and how pairs of simultaneously activated items can become jointly indexed and associated. Activating one pair member now also activates the other one indirectly via PFC with a short latency. We have further demonstrated that this kind of WM can readily be updated such that as new items are encoded, old ones are fading away whereby the active WM content is replaced.

Recall dynamics in the presented model are in most respects identical to our previous cortical associative memory models (Lansner 2009). Any activated memory item, whether randomly or specifically triggered, is subject to known and previously well characterized associative memory dynamics, such as pattern completion, rivalry, bursty reactivation dynamics, oscillations in different frequency bands, etc. (Lundqvist et al. 2010; Silverstein & Lansner 2011; Lundqvist et al. 2013; Herman et al. 2013). Moreover, sequential learning and recall could readily be incorporated (Tully et al. 2013). This could for example support encoding of sequences of items in WM rather than a set of unrelated items, resulting in reactivation dynamics reminiscent of e.g. the phonological loop (Baddeley et al. 1998; Burgess & Hitch 2006).

### The Case for Hebbian Plasticity

The underlying mechanism of our model is fast Hebbian plasticity, not only in the intrinsic PFC connectivity, but also in the projections from PFC to LTM stores. The former has some experimental support (Volianskis & Jensen 2003; Volianskis et al. 2015; Erickson et al. 2010; Park et al. 2014; Kauer et al. 2018) whereas the latter remains a prediction of the model. Dopamine D1 receptor (D1R) activation by dopamine (DA) is strongly implicated in reward learning and synaptic plasticity regulation in the basal ganglia (Wickens 2009). In analogy we propose that D1R activation is critically involved in the synaptic plasticity intrinsic to PFC and in projections to LTM stores, which would also explain the comparatively dense DA innervation of PFC and the prominent WM effects of PFC DA level manipulation (Arnsten & Jin 2014; Goto et al. 2010). In our model, the parameter *κ* represents the level of DA-D1R activation, which in turn regulates its synaptic plasticity. We typically increase kappa 4-8 fold temporarily in conjunction with stimulation of LTM and WM encoding, in a form of attentional gating. Larger modulation limits WM capacity to 1-2 items, while less modulation diminishes the strength of cell assemblies beyond what is necessary for reactivation and LTM maintenance.

When the synaptic plasticity WM hypothesis was first presented and evaluated, it was based on synaptic facilitation (Mongillo et al. 2008; Lundqvist et al. 2011). However, such non-Hebbian plasticity is only capable of less specific forms of memory. Activating a cell assembly, comprising a subset of neurons in an untrained STM network featuring such plasticity, would merely facilitate all outgoing synapses from active neurons. Likewise, an enhanced elevated resting potential resulting from intrinsic plasticity would make the targeted neurons more excitable. In either case, there would be no coordination of activity specifically within the stimulated cell assembly. Thus, if superimposed on an existing LTM, such forms of plasticity may well contribute to WM, but they are by themselves not capable of supporting encoding of novel memory items or the multi-modal association of already existing ones. In contrast, in our previous work (Fiebig & Lansner 2017) we showed that fast Hebbian plasticity similar to STP (Erickson et al. 2010) allows effective one-shot encoding of novel STM items. In the extended model proposed here, PFC can additionally bind and bring on-line existing but previously unassociated LTM items across multiple modalities by means of the same kind of plasticity in backprojections from PFC to parieto-temporal LTM stores.

On a side note, our implementation of fast Hebbian plasticity reproduces a remarkable aspect of STP or Labile LTP: it decays in an activity-dependent manner rather than with time (Volianskis & Jensen 2003; Volianskis et al. 2015; Kauer et al. 2018). Although we used the BCPNN learning rule to reproduce these effects, we expect that other Hebbian learning rules allowing for neuromodulated fast synaptic plasticity could give comparable results.

### Experimental support and Testable predictions

Our model has been built from available relevant microscopic data on neural and synaptic components as well as modular structure and connectivity of selected cortical areas in macaque monkey. The network so designed generates a well-organized macroscopic dynamic working memory function, which can be interpreted in terms of manifest behavior and validated against cognitive experiments and data. Our model provides a powerful tool to investigate and examine the link between microscopic and macroscopic level processes and data. It suggests novel mechanistic hypotheses and inspiration for planning and performing experiments that can develop further the model, or potentially falsify it.

Unfortunately, the detailed neural processes and dynamics of our new model are not easily accessible experimentally as they are intrinsically expressed at multiple scales, e.g. mesoscopic field potentials and population spiking at macroscopic spatial scales. In consequence, it is difficult to find direct and quantitative results to validate the model. Yet, in analyzing our resulting bottom-up and top-down delays we drew an analogy to a split-brain experiment (Tomita et al. 1999) because of its clean experimental design (even controlling for subcortical pathways) and found similar temporal dynamics in our highly subsampled cortical model. The timing of inter-area signals also constitutes a testable prediction for multi-modal memory experiments. Furthermore, reviews of intracranial as well as electroencephalography (EEG) recordings conclude that theta band oscillations play an important role in long-range communication during successful memory retrieval (Johnson & Knight 2015; Sauseng et al. 2004). With respect to theta band oscillations in our model, we have shown that STM leads the LTM networks during maintenance, engages bi-directionally during recall (due to the STM-LTM loop), and lags during stimulus-driven encoding and LTM activation, reflecting experimental observations (Anderson et al. 2010). These effects are explained by our model architecture, which imposes delays due to the spatial extent of networks and their distances from each other. Fast oscillations in the broad gamma band, often nested in the theta cycle, are strongly linked to local processing and activated memory items in our model, also matching experimental findings (Canolty & Knight 2010; Johnson & Knight 2015). Local frequency coupling is abundant with significant phase-amplitude coupling (e.g. Figure 3B), and was well characterized in related models (Herman et al. 2013).

The most critical requirement and thus prediction of our theory and model is the presence of fast Hebbian plasticity in the PFC backprojections to parieto-temporal memory areas. Without such plasticity, our model cannot explain the necessary STM-LTM binding. This plasticity is likely to be subject to neuromodulatory control, presumably with DA and D1R activation involvement. Since STP decays with activity, a high noise level could be an issue since it could shorten WM duration (see *The Case for Hebbian Plasticity*). The evaluation of this requirement is hampered by little experimental evidence and a general lack of experimental characterization of the synaptic plasticity in long-range corticocortical projections.

One of the neurodynamical manifestations of the fast associative plasticity in the PFC backprojections is a functional coupling between LTM stores. Importantly, this long-range coupling in our model is mediated by the PFC network alone, as manifested during delay period free of any external cues, and is reflected in the synchronization of fast gamma oscillations. Although the predominant view has been that gamma is restricted to short distances, there is growing evidence for cortical long-distance gamma phase synchrony between task-relevant areas as a correlate of cognitive processes (Tallon-Baudry et al. 1998; Doesburg et al. 2008) including WM (Palva et al. 2010). In this regard, our model generates even a more specific prediction about the notable temporal enhancement of gamma phase coupling over the delay period, which could be tested with macroscopic human brain recordings, e.g. EEG or magnetoencephalography (MEG), provided that a WM task involves a sufficiently long delay period.

Finally, our model suggests the occurrence of a double peak of frontal network activation in executive control of multi-modal LTM association (see STM population activity during WM Maintenance in Figure 5). The first one originates from the top-down control signal itself, and the second one is a result of corticocortical reentry and a successful activation of one or more associated items in LTM. As such, the second peak should also be correlated with successful memory maintenance or associative recall.

Furthermore, our model also makes specific predictions of neuroanatomical nature about the density of corticocortical long-range connectivity. For example, as few as six active synapses (*Methods*) onto each coding pyramidal neuron are sufficient to transfer specific memory identities across the cortical hierarchy and to support maintenance and recall.

### Role of fast Hebbian plasticity in Variable Binding

The “binding problem” is a classical and extensively studied problem in perceptual and cognitive neuroscience, see e.g. Zimmer et al. (2012). Binding occurs in different forms and at different levels, from lower perceptual to higher cognitive processes (Reynolds & Desimone 1999; Zimmer et al. 2006). At least in the latter case, WM and PFC feature quite prominently (Cer & O’Reily 2012) and this is where our WM model may provide further insight.

Variable binding is a special case and a cognitive kind of neural binding in the form of a variable – value association of items previously not connected by earlier experience and learning (Cer & O’Reily 2012; Garnelo & Shanahan 2019). A simple special case is the association of a mathematical variable and its value “The value of x is 2”, i.e. x = 2. More generally, an object and a name property can be bound like in “Charlie is my parrot” such that <name> = “Charlie” (Figure 9). This and other more advanced forms of neural binding are assumed to underlie complex functions in human cognition including logical reasoning and planning (Pinkas et al. 2012), but have been a challenge to explain by neural network models of the brain (Legenstein et al. 2016; van der Velde & de Kamps 2015).

**Figure 9.**
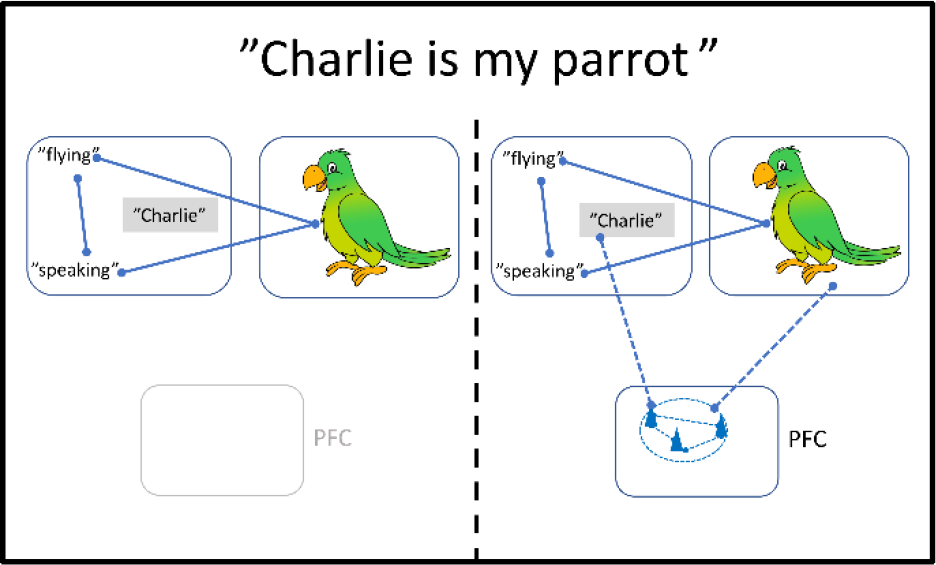
Variable-value binding via PFC. Initially the representation of “parrot” exists in LTM comprising symbolic and sub-symbolic components. When it is for the first time stated that “Charlie is my parrot”, the name “Charlie” is bound reciprocally by fast Hebbian plasticity via PFC to the parrot representation, thus temporarily extending the composite “parrot” cell assembly. Pattern completion now allows “Charlie” to trigger the entire assembly and “flying” or the sight of Charlie to trigger “Charlie”. If important enough or repeated a couple of times this association could consolidate in LTM.

Based on our WM model, we propose that fast Hebbian plasticity provides a neural mechanism that mediates such variable binding. The joint index to LTM areas formed in PFC/STM during presentation of a name – image stimulus pair serves to bind the corresponding LTM stored variable and value representations in a specific manner that avoids mixing them up. Turning to Figure 5 above, imagine that one of the LTMa patterns represent the image of my parrot and one pattern in LTMb, now a cortical language area, represents his name “Charlie”. When this and two other image – name pairs are presented they are each associated via specific joint PFC indices. Thereafter “Charlie” will trigger the visual object representation of a parrot, and showing a picture of Charlie will trigger the name “Charlie” with a dynamics as shown in the right-most panels of Figure 5. Here as well, flexible updating of the PFC index will avoid confusion even if in the next moment my neighbor shouts “Charlie” to call his dog, also named Charlie.

Recent experiments have provided support for the involvement of PFC in such memory related forms of feature binding (Zmigrod et al. 2014). Gamma band oscillations, frequently implicated when binding is observed, are also a prominent output of our model (Tallon-Baudry & Bertrand 1999). Work is in progress to uncover how such variable binding mechanisms can be used in neuro-inspired models of more advanced human logical reasoning (Pinkas et al. 2013).

## Conclusions

We have formulated a novel indexing theory for WM and tested it by means of computer simulations, which demonstrated the versatile WM properties of a large-scale spiking neural network model implementing key aspects of the theory. Our model provides a new mechanistic understanding of the targeted WM and variable binding phenomena, which connects microscopic neural processes with macroscopic observations and cognitive functions in a way that only computational models can do. While we designed and constrained this model based on macaque data, the theory itself is quite general and we expect our findings to apply also to mammals including humans, commensurate with changes in key model parameters (cortical distances, axonal conductance speeds, etc.). Many aspects of WM function remains to be tested and incorporated, e.g. its close interactions with basal ganglia (O’Reilly & Frank 2006).

WM dysfunction has an outsized impact on mental health, intelligence, and quality of life. Progress in mechanistic understanding of its function and dysfunction is therefore very important for society. We hope that our theoretical and computational work provides inspiration for experimentalists to scrutinize the theory and model, especially with respect to neuromodulated fast Hebbian synaptic plasticity and large-scale network architecture and dynamics. Only in this way can we get closer to a more solid understanding and theory of WM, and position future computational research appropriately even in the clinical and pharmaceutical realm.

## Supplementary Information

### Model Robustness

Our model incorporates a plethora of biological constraints, such as estimates on the extent and distance of areas (e.g. STM patch size approximates macaque dlPFC, and is 40mm from ITC), laminar cell distributions 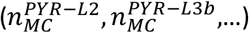, hypercolumnar size, etc. The model also abides by various electrophysiological constraints, such as plausible EPSP, IPSP sizes, estimates on laminar connection densities, characterization of cortical FF/FB pathways, estimates on axonal conductance speeds, dendritic arbor sizes (branching factors), commonly accepted synaptic time-constants for various receptor types, depression, adaptation, and builds on top of established models we adapted, such as the neuron model or the synaptic resource model. References to many of these constraints can be found throughout the Method Section.

Because our model is quite complex and synthesizes many different components and processes it is beyond the scope of this work to perform a detailed parameter sensitivity analysis. However, from our extensive simulations we conclude that it is robust and degrades gracefully. Almost all uncertain parameters can be varied ±30% without breaking WM function. The model is dramatically subsampled and scaling up would be possible. This could be expected to further improve overall robustness. Highly related modular cortical network models have been studied extensively elsewhere(Lundqvist et al. 2010; Tully et al. 2013; Lundqvist et al. 2011; Fiebig & Lansner 2017; Tully et al. 2014), so here we prioritize novel aspects, namely the parameterization of corticocortical connectivity and spatial scale.

In the feedback pathway, a mere 0.6% connectivity is sufficient to support LTM activation in maintenance and recall. As rigorous testing (not shown here) revealed, lower connectivity degrades WM capacity, unless we increase the total number of co-active STM cells by other means. Forward connectivity can be even lower (0.015% in this model), because terminal clusters in STM are smaller and provide more information contrast (*Corticocortical Connectivity*). In both cases, our model uses these low density values, but they could be increased or decreased if single synaptic currents are reduced/increased respectively. Somewhat peculiarly, we also found that we needed to increase the corticocortical conductance of the backprojections (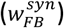) by the same factor 1.8 (over the local conductance gain 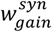) as another detailed model account of macaque visual cortex(Schmidt et al. 2015) to achieve functional WM at the stated long-distance connection probabilities.

There is an upper, but no lower limit on corticocortical distances in our model. When conduction delays exceed 65 ms (130 mm), STM feedback can no longer activate the LTM network, because bursts desynchronize before they arrive. On the other hand, STM and LTM could even be adjacent as we briefly mentioned at the end of the result section. Additionally, there is a minimum spatial scale to each component network. If we reduce the spatial extent (and thus the connection delays between HCs) by 45%, theta-like oscillations degrade and break at 20%, when the largest inter-HC delays fall below 5 ms. Spiking activity of activated memories collapses into a single brief burst (Figure 3 – Supplement 2, cf. Figure 3D), which degrades learning and effective information transmission both within and across networks. Networks may be much smaller however, if this is compensated by slower axonal conductance velocities (<2 mm/ms).

### Supplementary Figures+legends

**Figure 3 – Supplement 1.**
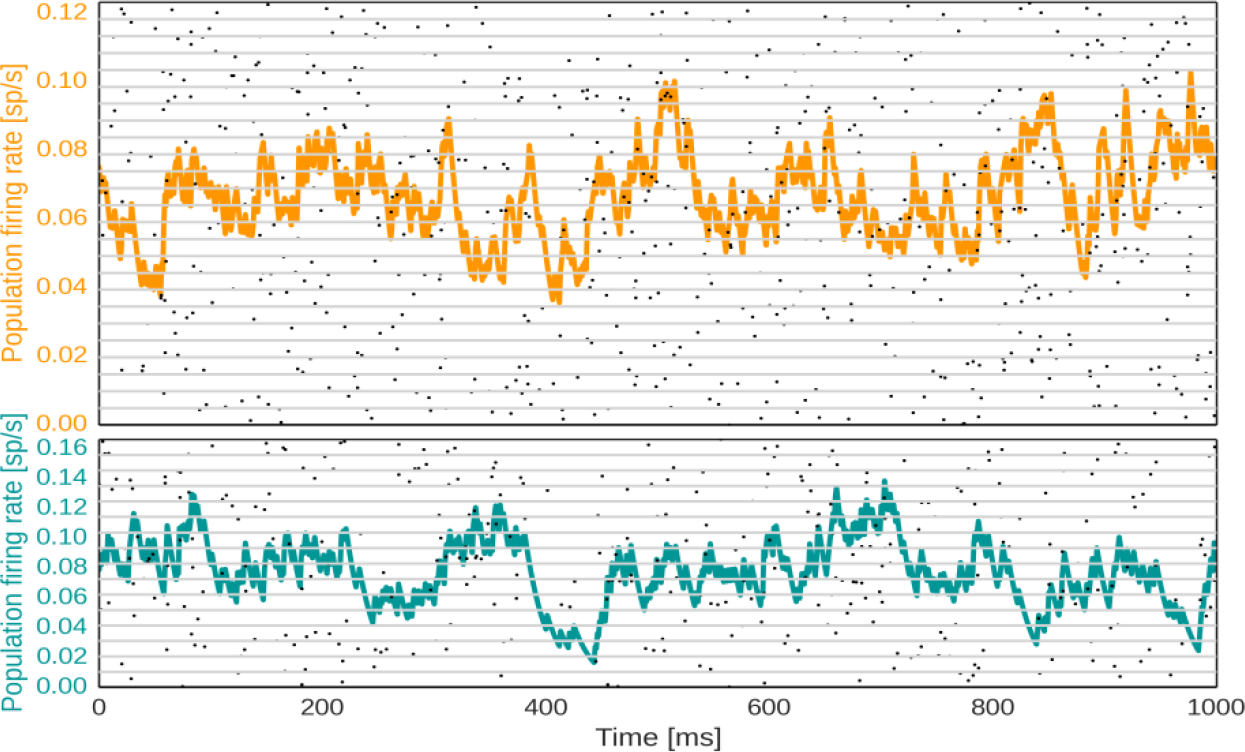
Basic Network behavior in spike rasters and population firing rates under low input. The untrained networks STM (top) and LTM (bottom) feature low rate, asynchronous activity (CV2 = 0.7±0.2). The underlying spike raster shows layer 2/3 activity in each HC (separated by grey horizontal lines) in the simulated network.

**Figure 3 – Supplement 2.**
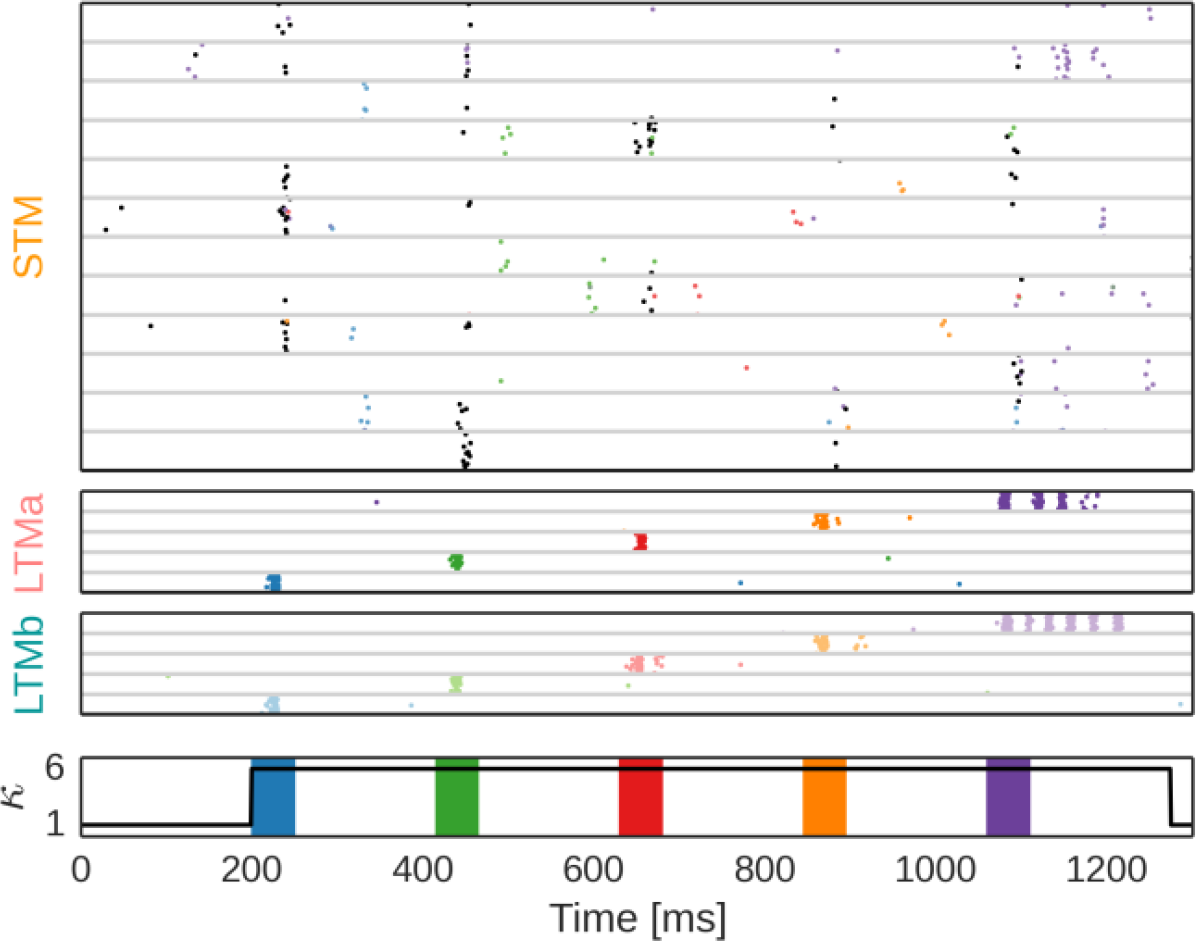
Network activity during plasticity-modulated stimulation with 20% spatial extent. Subsampled spike raster of the layer 2/3 population in a Hypercolumn of STM (top), and five coding minicolumns in LTMa (2^nd^ row) and LTMb (3^rd^ row) respectively during plasticity-modulated stimulation (i.e. encoding) of five paired LTM patterns. Without sufficient conduction delays, memory activations collapse into very brief bursts (with the exception of the last pattern here) and STM cannot effectively activate from or subsequently encode such brief activations (cf. Figure 2D, and **Supplementary Figure 6**).

**Figure 4 – Supplement 1.**
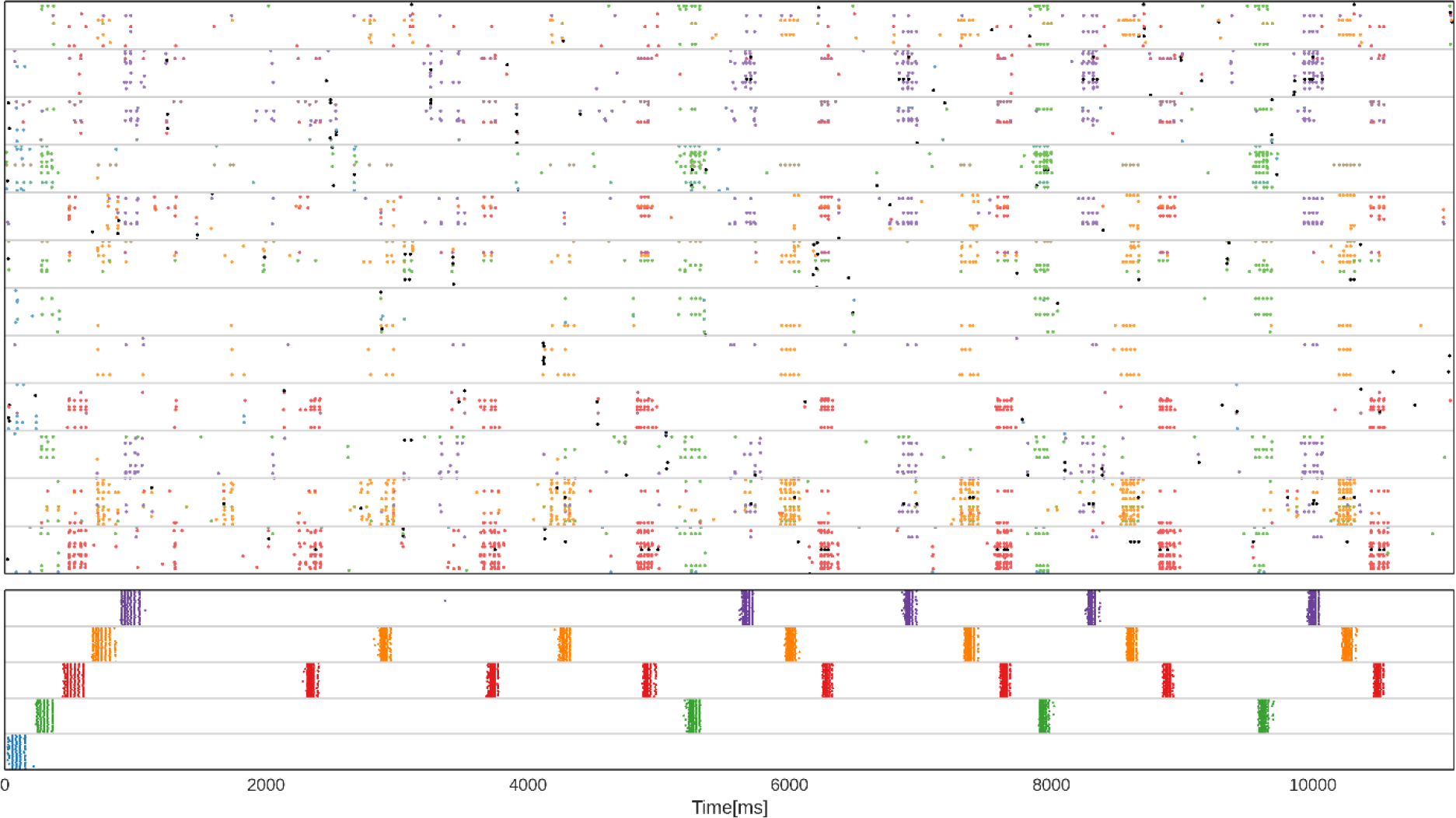
Encoding and feedback-driven reactivation of long-term memories. Subsampled spike raster of STM (top) and LTM (bottom) during encoding and subsequent maintenance of five memories (the first pattern is not maintained in this simulation). During the initial plasticity-modulated stimulation phase, five LTM memories are cued via targeted 50 ms stimuli (shown underneath). Plasticity of STM and its backprojections is modulated during this initial memory activation (cf. Figure3D). Thereafter, a strong noise drive to STM causes spontaneous activations and plasticity-induced consolidation of pattern-specific subpopulations in STM. Backprojections reactivate associated LTM memories. **Top:** STM spike raster shows layer 2/3 activity in a single HC. MCs are separated by grey horizontal lines. STM spikes are colored according to each cell’s dominant LTM pattern-correlation, similar to Figure 2D. **Bottom:** LTM spike raster only shows the activity of five coding MC in a single LTM HC, but indicates the activation of distributed LTM memory patterns. LTM spikes are colored according to the pattern-specificity of each cell.

**Figure 4 – Supplement 2.**
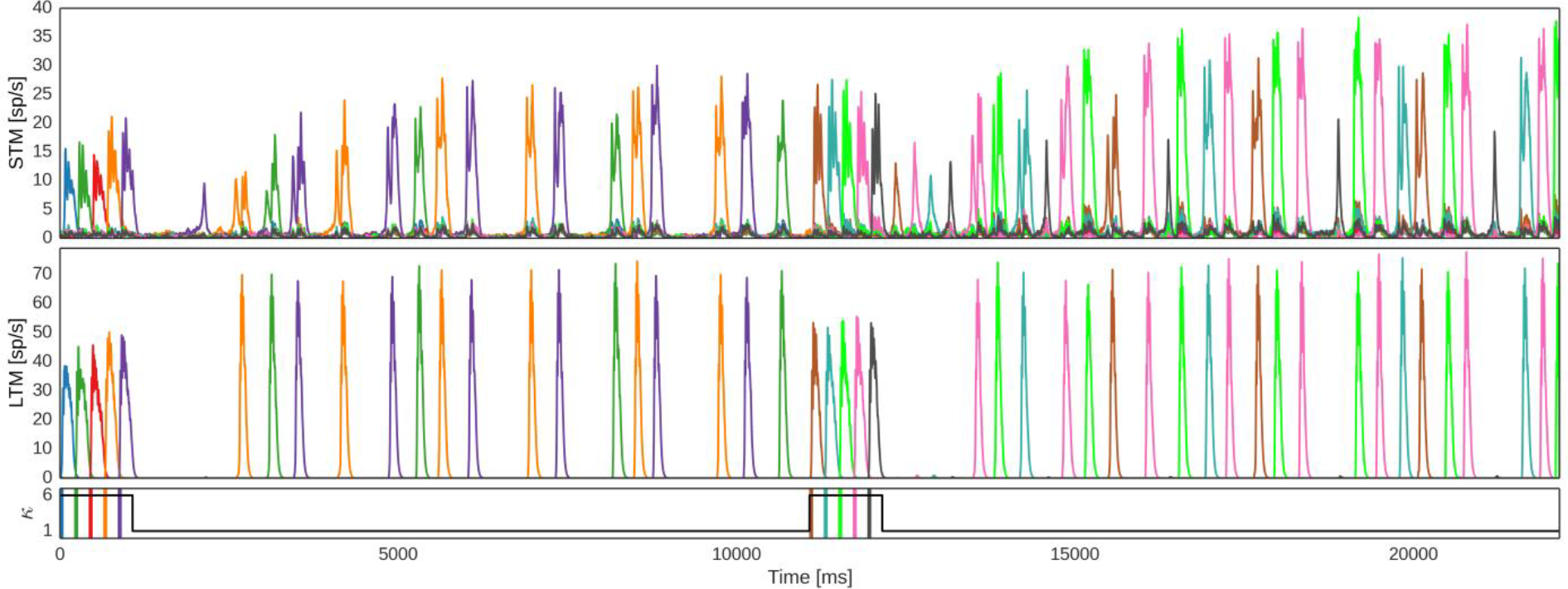
Spike rates during WM updating. Population firing rates of pattern-specific subpopulations in STM and LTM during encoding and subsequent maintenance of two sets of five LTM memories. After encoding and 10 s maintenance of the first set, WM contents are overwritten with the second set of memories, maintained thereafter in spontaneous reactivation events. Bottom: Stimuli to LTM and modulation of plasticity.

**Figure 4 – Supplement 3.**
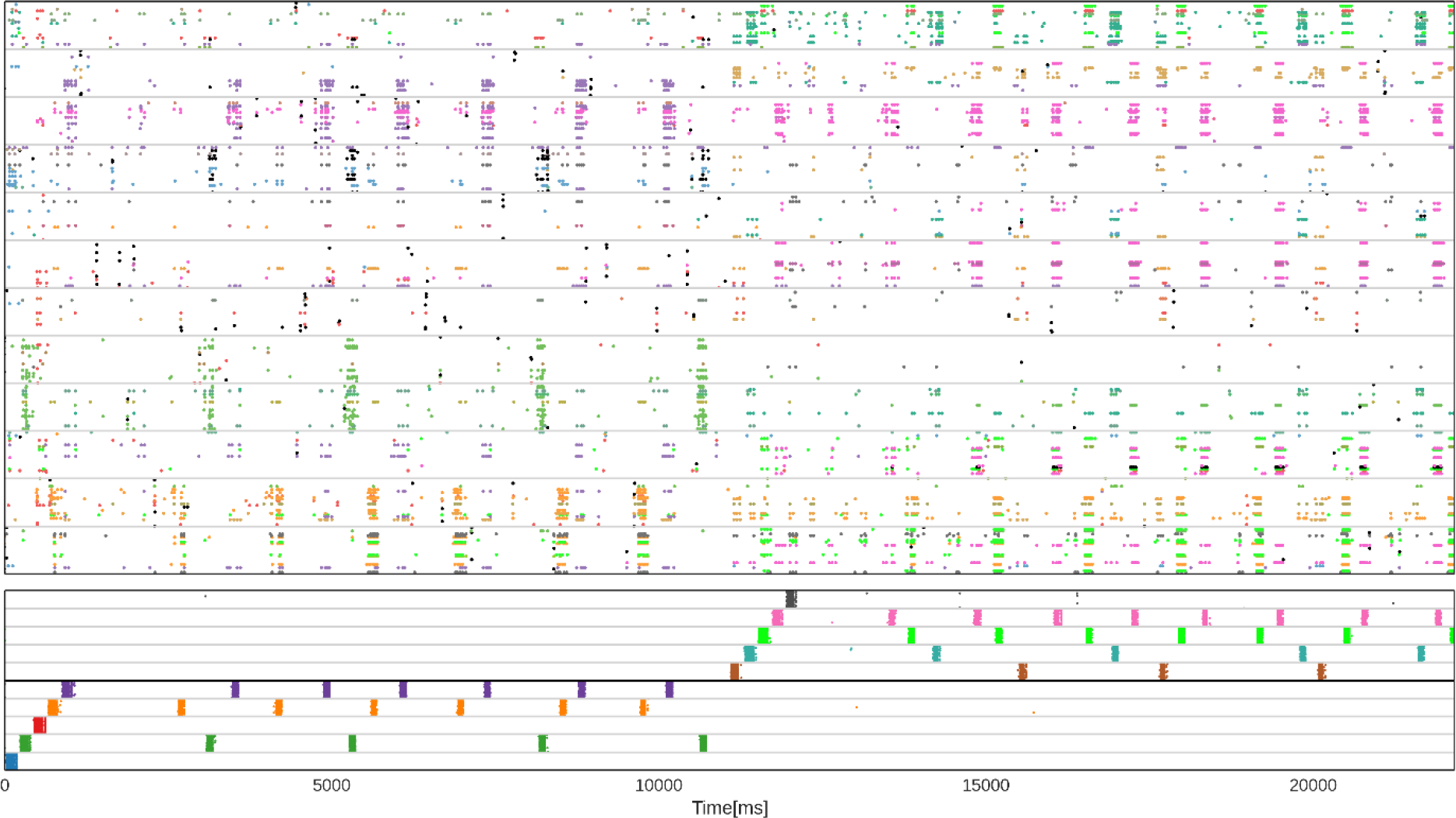
Spike raster during WM updating. Subsampled spike raster of the layer 2/3 population in a Hypercolumn of STM (top) and LTM (bottom) respectively during encoding and subsequent maintenance of two sets of five LTM memories. STM spikes are colored according to each cells dominant pattern-selectivity. LTM spikes are colored according to the pattern-specificity of each cell. After encoding and 10 s maintenance of the first set, WM contents are overwritten with the second set of memories, maintained thereafter. Plasticity is temporarily boosted during the initial activation of LTM attractors (see preceding figure). Strong noise drive to STM causes spontaneous reactivations and consolidation of pattern-specific subpopulations in STM following each stimulation period.

**Figure 7 – Supplement 1.**
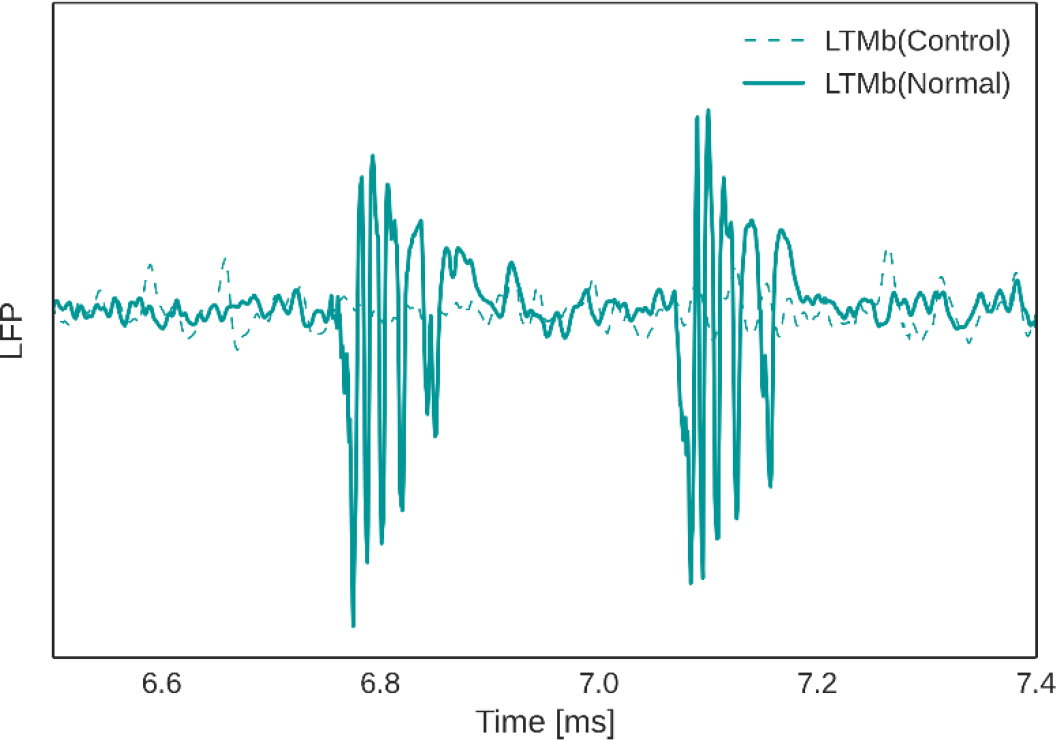
Exemplary recording of the Local Field Potential (LFP) signal in LTMb following two cued activations of LTMa after learning and maintenance of associative LTMa-LTMb memory pairs (normal) or non-associative LTMa memories without concurrent LTMb activation (control). While the LFP signal shows clear activation of associated LTMb items, LTMa specific cues do not elicit memory activations in LTMb in the control case.

**Figure 5 – Supplement 1.**
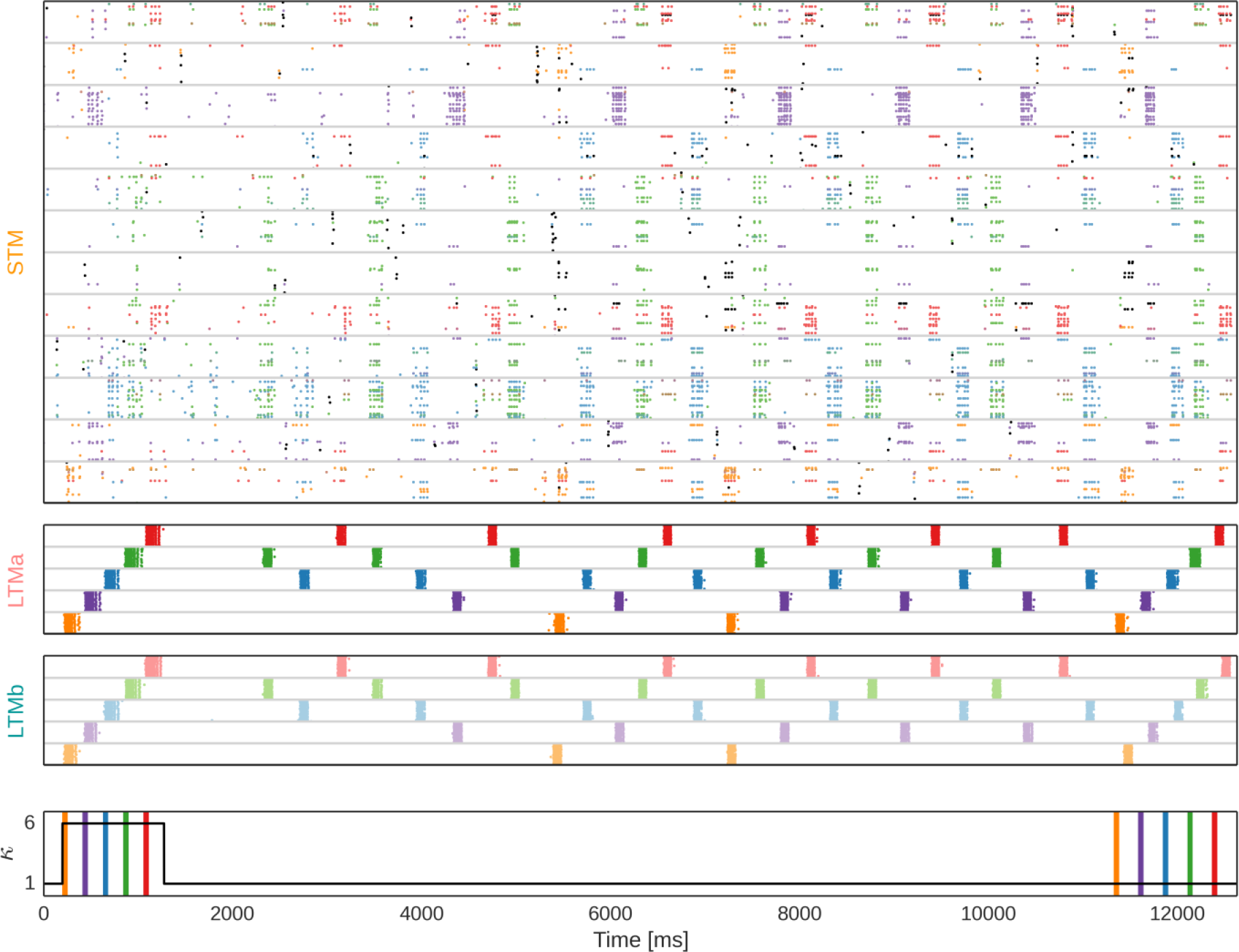
Spiking activity in the three networks, during the multi-modal LTM binding task. Subsampled spike raster of the layer 2/3 population in a Hypercolumn of STM (top), and five coding minicolumns in LTMa (2^nd^ row) and LTMb (3^rd^ row) respectively during plasticity-modulated stimulation (i.e. encoding), subsequent maintenance, and associative cued recall of five paired LTM patterns (orange,purple,blue,green,red). Minicolumns are separated by grey horizontal lines. STM spikes are colored according to each cells dominant memory pair-selectivity. LTM Spikes are colored according to the memory pair-specificity of each cell in slightly shifted hues to illustrate that LTMa and LTMb code for different, but associated memories. Bottom: Stimuli to LTM and modulation of plasticity. Note the cued recall of all five memories at the end.

## Acknowledgements

This work was supported by the EuroSPIN Erasmus Mundus doctoral program, SeRC (Swedish e-science Research Center), and StratNeuro (Strategic Area Neuroscience at Karolinska Institutet, Umeå University and KTH Royal Institute of Technology). The simulations were performed using computing resources provided by the Swedish National Infrastructure for Computing (SNIC) at PDC Centre for High Performance Computing. We are grateful for helpful comments and suggestions from Drs Jeanette Hellgren Kotaleski, and Arvind Kumar.

